# Nanoprotrusion enlargement conducts cytocapsular tube elongation along the path of least resistance for metastasis initiation

**DOI:** 10.1101/2025.02.10.637526

**Authors:** Tingfang Yi, Gerhard Wagner

## Abstract

Cancer metastasis is a major source of cancer lethality. Recently, we reported that the ATP-dependent calcium pump PMCA2 is enriched in cytocapsular membranes as biomarker of native malignant tumors. Cytocapsular tubes (CCTs) provide membrane-enclosed freeways for protected and bi-directional cancer metastasis in cytocapsular tumor network systems (CTNSs). It is obscure how CCTs initiate and elongate in heterogeneous human tissues and organs against diverse blocks of resistance. Here, we report that thin CCT nanoprotrusions (NPs) enlarge in directions of least resistance to grow into enlarged NPs. Subsequently, enlarged NPs develop into initial CCTs (ICCTs) followed by development into full CCTs. The full CCTs elongate in paths of least resistance over long distances, as observed in 34 kinds of human tissues and organs. Sideways CCT branching proceeds via a NP bifid format, which increases CCT numbers and expands 3D CCT networks in tissues. CCT regeneration additionally drives repeated cancer metastasis. CCT superstructures facilitate full cancer metastasis. This study demonstrated nanoprotrusion-conducted CCT elongation along the path of least resistance and branching morphogenesis in bifid branching style. The cytocapsular membrane system expansion mechanics of CCTs and CCT networks described here may open new avenues for native cancer research and CTNS-targeted effective cancer cure. (200 words)

**Significance statement:** The recent identification of PMCA2 as a biomarker for malignant tumors, allows imaging very early native cancer growths on culture plates *in vitro* and in human bodies as revealed from snapshot images of numerous cancer biopsies *in vivo*. Aggressive cancer metastasis, cancer progression, repeated cancer relapse, limited cancer therapy outcomes, pan-cancer drug resistance, immune cell attack escape, and immunotherapy non-responsive “cold” tumors are unmet challenges in clinical cancer therapy. However, the cytocapsular tube (CCT) elongation mechanisms in human tissues/organs with heterogeneous resistances are still unclear. Here, we report that CCT nanoprotrusions (NPs) extend into enlarged NPs apparently towards tissue of least resistance and subsequently develop into initial CCTs. Intracytocapsular oncocells released cytocapsulasomes to fuse into the membranes of initial CCTs and increase cytocapsular membrane areas. This enables oncocells to push and deform initial CCT membranes and to generate new CCTs with new nanoprotrusion layers. Furthermore, CCT branching morphogenesis develops in a bifid format, significantly increasing CCT numbers and expanding 3D CCT networks in all kinds of human tissues. CCT superstructures promote complete cancer metastasis in AMCC complexes. Repeated CCT regeneration drives repeated cancer relapses. This study reveals the CCT elongation laws, and elucidates that CCT elongation and branching morphogenesis drives cytocapsular membrane protected cancer metastasis, and that repeated CCT regeneration promotes repeated tumor relapses *in vivo*.

## Introduction

Recently, we identified the biomarker membrane protein ATP-dependent calcium pump PMCA2-enriched cytocapsular membranes as hallmarks of native malignant tumors ^1-3^. Cancer metastasis is the major source of cancer deaths^4-12^. New insights into mechanisms of metastases have come from the discovery of cytocapsular tubes (CCTs) which provide the major pathways for native cancer dissemination^1-3^. CCT membranes extend and generate many nanoprotrusions anchoring CCTs in place^1^. Human tissues and organs in cancer micro-environments harbor extremely heterogeneous materials/factors, including all kinds of cells, extracellular matrix (ECM), cell-cell and cell-ECM interactions, solid stresses, interstitial fluid pressure, physical microarchitectures, which are resistances of CCT elongation^13-26^. It is obscure how CCTs elongate and expand in heterogeneous human tissues and organs.

Here, we investigated CCT elongation in cancers in 34 kinds of human tissues and organs, and found that: (i) CCT nanoprotrusion enlargement conducts CCT elongation along the path with least resistance, (ii) CCT bifid branching morphogenesis increases CCT numbers and expands CCT networks. (iii) CCT regeneration in AMCC associates with new cycles of cancer metastasis, and (iv) CCT superstructures promote complete cancer metastasis. These discoveries may facilitate native cancer researches, cancer drug research and development, and clinical cancer applications.

## Results

### Cytocapsular nanoprotrusions enlarge and generate initial cytocapsular tubes followed by new CCT formation and CCT elongation

At 24hour (h) after implanting into CC/CCT culture kits, single Bxpc3 pancreas cancer cells generate cytocapsulas (CCs) with many nanoprotrusions (NPs) and NP bunches, which interconnect and form NP networks surrounding CCs (**Extended Data Fig. 1A**, n = 316 tested cytocapsular oncocells). When ecellulated CCs (ECCs) of single cytocapsular oncocells appear they also exhibit NPs and NP bunches but degrade and disappear after about 6 hrs. (**Extended Data Fig. 1B**, n=56 ECCs). These results show that single cytocapsular oncocells’ cytocapsular membranes extend and generate many NPs and NP bunches, which interconnect and form NP networks surrounding cytocapsular oncocells (**Extended Data Figs. 1A-1B**).

At 36h, Bxpc3 pancreas intracytocapsular oncocells proliferate and generate cytocapsular tumorspheres wrapped in the enlarged CCs, which engender larger NP networks surrounding enlarged CCs (**Extended Data Fig. 1C**, n= 98 cytocapsular tumorspheres). Transmission electron microscope (TEM) analyses of breast cancer stem cell (BCSC) HMLER (CD44^high^/CD24^low^)^FA^ cell NPs show that the NPs are coated with glycocalyx (a mixture of glycoproteins and glycolipids) in variable thickness (up to 78nm in thickness, **Extended Data Fig. 1D**). At 36h, the NP sizes are variable (21nm ∼ 82nm in diameter/width, 300-2700nm in length). Further TEM analyses of these BCSC cytocapsular oncocell NP bunches show that two or more straight or curved NPs align together tightly/loosely and form NP bunches, coated with glycocalyx layers in wider thickness range (2nm∼1000nm in thickness, **Extended Data Fig. 1E**). In NP bunches, single NP sizes are 18nm ∼ 32nm in diameter/width. The NP bunch sizes are variable (40nm ∼ 152nm in width, **Extended Data Fig. 1E**). NP generate branches, indicating NP branching morphogenesis is a resource of NP number increase and NP network formation surrounding CCs (**Extended Data Fig. 1E**). Glycocalyx may cause the blur/smear effects of NP networks in bright field and immunohistochemistry fluorescence microscope imaging analyses.

Subsequently, at 48h, when observing the larger tumorspheres which also have NPs, one or a few single Bxpc3 cytocapsular tumorspheres’ cytocapsular NPs elongate in length (up to 100μm in length, n=46 NPs) and enlarge in size (up to 0.1μm in diameter/width, n=46 enlarged NPs), and link CCs of cytocapsular tumorspheres (CTs, **Extended Data Fig. 2A**). There is no oncocell in migration in these early enlarged NPs (82nm ∼ 0.1μm in diameter/width) that directly link two CTs (**Extended Data Fig. 2A**, n=126 enlarged NPs). Not all NPs have equal chances to enlarge, and only the shortest NPs that can reach nearby CTs are preferred to grow up and enlarge (**Extended Data Fig. 2A**, n=126 enlarged NPs). This indicates that NP networks sense distances between CTs, and the NPs with shortest distance are preferred to grow up and enlarge in sizes.

At 56h *in vitro*, single Bxpc3 cytocapsular tumorsphere-enlarged NPs that link CTs continue to grow. They generate cytocapsular tubes (CCTs) of various diameters/widths from 0.1μm to ∼ 3μm (**Extended Data Fig. 2B**, n=65 NPs). In the NP enlargement-generated early CCTs with sizes of only 0.1μm ∼ 0.5μm in diameter/width, there are oncocells in migration inside, indicating that these early CCTs have biological functions for CCT-conduced oncocell migration inside. These NP enlargement-engendered early CCTs are termed initial CCTs (ICCTs). Established CCTs with sizes of >0.5μm in diameter/width (**Extended Data Fig. 2C**) allow bi-directional oncocell migration between cytocapsular tumorspheres *in vitro*. There are many NPs surrounding ICCTs and CCTs, anchoring ICCTs and CCTs in place. The above data demonstrated that a few cytocapsular NPs with shortest distance to nearby CTs and least resistance of molecular communications between CTs are selectively growing into ICCTs and CCTs for oncocell migration inside *in vitro*. NP enlargement and CCT development are terminated in ecellulated cytocapsulas (from cytocapsular oncocells/tumorspheres, **Extended Data Figs. 1 and 2**), demonstrating that NP enlargement, ICCT development and CCT elongation are dependent on incytocapsular oncocell driving (cytocapsulasomes and other molecules).

Consistently, in the nuperphase tumors^2^ of invasive breast carcinomas, there are many initial CCTs *in vivo* (ICCTs, 0.2μm ∼ 0.5μm in diameter/width, 10μm ∼ 230μm in length in sectioned specimens, n=215 ICCTs, from 112 breast cancer patient; Fig. S3A). There are oncocells in migration in some initial CCTs (**Extended Data Fig.S3A**). These observations of initial CCTs in clinical invasive breast carcinoma tissues evidenced that clinical breast cancer cytocapsular nanoprotrusions (NPs) grow into initial CCTs in tissues *in vivo*. The initial CCTs elongate in tiny pores or small cavities with least resistance (**Extended Data Fig.S3A**), indicating the NPs sense resistance in tissues, and NPs with least resistances grow up, enlarge and engender initial CCTs. Next, we examined initial CCTs in cancers of 34 kinds of human tissues/organs (with 289 subtypes). Indeed, initial CCTs are presented in all the investigated cancers of 34 kinds of human tissues/organs (289 subtypes of cancers, **Extended Data Fig. 4**). In some instances, the sectioned thin initial CCT fragments are straight and crossing tissues. Most sectioned initial CCT fragments are in curved or irregular morphologies in tissues. Later, the breast cancer initial CCTs grow up into CCTs (0.8μm ∼ 6μm in diameter/width) and with oncocells in dissemination inside. Both initial CCTs and CCTs are anchored in local positions in tissues, and CCT morphologies and locations largely reflect the initial CCTs’ morphologies and locations from which they grow up in tissues (**Extended Data Fig. 3A-B**). In summary, cytocapsular NPs grow into CCTs for oncocell migration and cancer metastasis *in vitro* and in tissues *in vivo*, and CCTs anchored in place in tissues reflect/record the locations and paths of initial CCTs and the selected NPs (**Extended Data Fig.3C**).

### Cytocapsular tubes elongate in the paths with least resistance

Next, we investigated elongation of super-long CCTs in tissues *in vivo*. In some instances, single pancreas adenocarcinoma cytocapsular oncocells generate very long, bent and curved CCTs (>7,000μm in length in sectioned cancer tissue specimens, >700-folds of average single cell diameter of 10μm) in the same plane in heterogeneous tissues, and being contained in a macro bent space (**Figs. 1A-B**). The super-long, super-large, oval-shaped and bent CCT suggests that the long pancreas cancer CCT elongates in a path with least resistance, which is sandwiched and bent by two layers of substantial resistances (the above big oval-shaped CCT bunch belt and the below big oval-shaped dense CCT strand belt, **Fig 1A**). The super-large, oval-shaped, bent CCT superstructure harbors numerous small, near periodic or aperiodic waves, or irregular curves. At the crest and trough positions, CCTs meet physical resistances from extracytocapsular tissues, and choose to change CCT elongation directions and extend along the paths with least resistance. In near periodic waves, the average wave heights are 52 μm ± 6.2 μm, and average wavelengths (λ) are 61 μm ± 3.4 μm (n=122 near periodic waves in tissues). These evidences demonstrated that the CCT experienced variable resistance at micro local position and always elongate forward in a path with least resistance (**Fig.1B**).

**Fig. 1.**
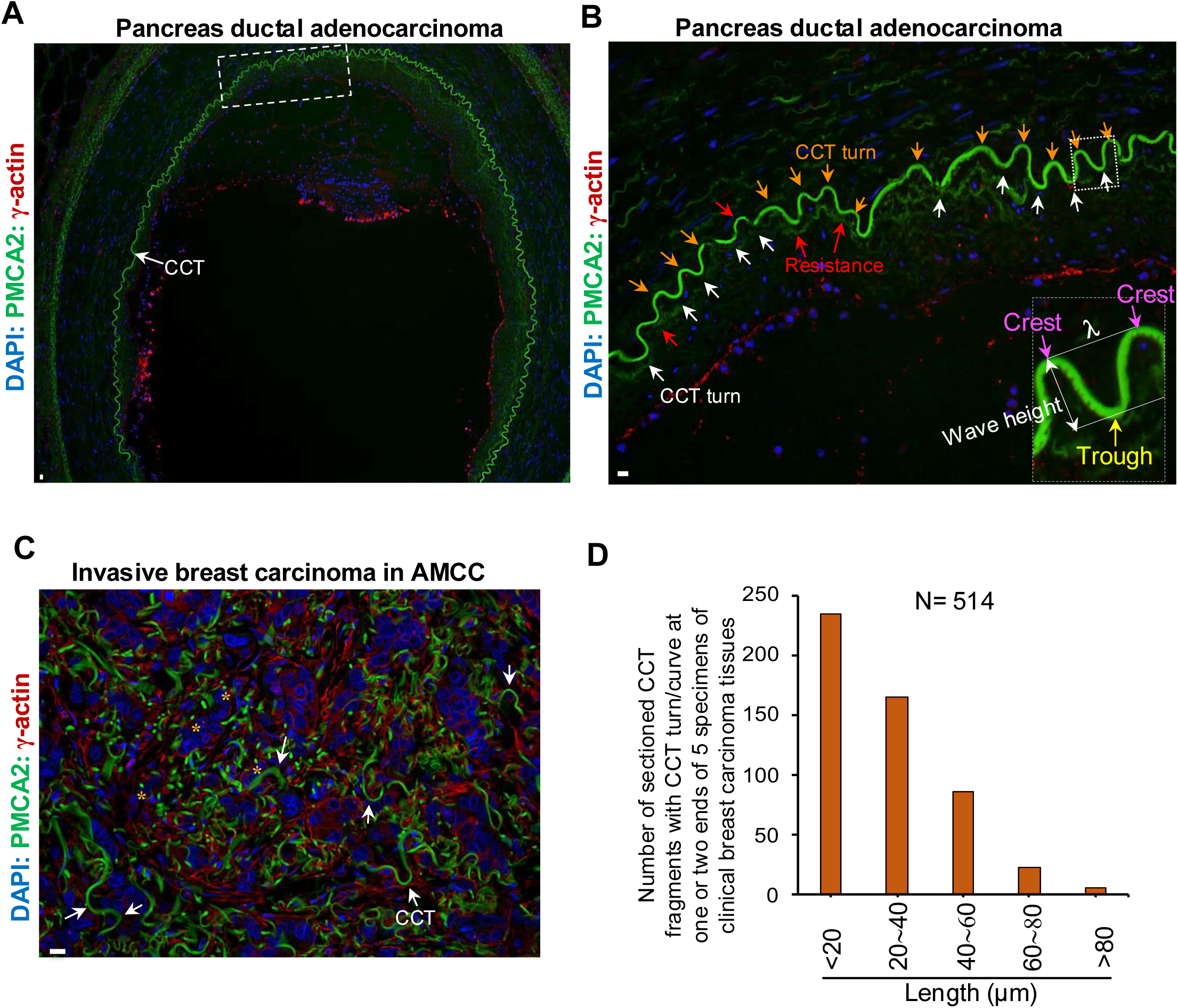
Cytocapsular tubes elongate along the path of least resistances in tissues *in vivo*. (A) Representative immunohistochemistry (IHC) fluorescence microscope image of a single pancreas cytocapsular oncocell generated, a part of a single, super-long CCT with many short waves, irregular curves and super-large CCT superstructure in a superlarge, bent, oval-shaped morphology in heterogeneous pancreas ductal adenocarcinoma tissue in the same plane. The macro resistances from the above and lower dense tissues force the super-long pancreas CCT to form a bent, oval-shaped superlarge morphology. The framed area is enlarged and shown in (B). CCT (white arrow) is shown. (**B**) An enlarged CCT fragment from (A). This CCT fragment has many CCT near periodic or aperiodic waves and irregular curves with short wave/curve height. Each pancreas CCT turn (orange arrows show crest/upper turns, white arrows show trough/bottom turns) is caused by the local micro resistances (red arrows) from the dense tissues along the pancreas CCT elongation direction. Then, the pancreas CCT nanoprotrusions (NPs) with least resistance enlarge and form pancreas enlarged NPs, followed by pancreas initial CCT generation in the least resistance, and established pancreas CCT formation in the least resistance. Therefore, form a pancreas CCT turn. An inserted enlarged near CCT wave panel shows the crest, trough, wave height and wave length (λ) of CCT near wave. (**C**) Representative IHC fluorescence microscope image of invasive breast carcinoma in acytocapsular mass-CC/CCT complex (AMCC) with many sectioned CCT fragments with CCT turns/curves. (**D**) Quantitative statistical analysis of number of sectioned CCT fragments with CCT turns/curves of clinical breast carcinoma tissues. Scale bar, 10μm.

Furthermore, single pancreas adenocarcinoma cytocapsular oncocell generate super-long, (>8,200 μm in length in sectioned cancer tissues >820-folds of average single cell diameter of 10μm), super-large, bent, flat ring-shaped CCT in the same plane in heterogeneous tissues (**Extended Data Fig. 5**). This long pancreas cancer CCT elongates in a path with least resistance sandwiched and bent by two layers of substantial resistances (the above super-large, flat ring-shaped and dense tissue cell layer and the below big flat ring-shaped dense CCT strand belt, **Extended Data Fig. 5**). Moreover, single prostate adenocarcinoma cytocapsular oncocells generate super-long, (>20,500μm in length in sectioned cancer tissues, >2,050-folds of average single cell diameter of 10μm), super-large, irregular-shaped, continuous, and highly bent CCT in superstructures in the same plane in heterogeneous tissues (**Extended Data Fig. 6**). The very-long single prostate cancer CCT with many small or big curves consistently elongates along the path with least resistance in the heterogeneous tissues (**Extended Data Fig. 6**). In colon adenocarcinoma tissues, single colon cytocapsular oncocells generate very long, highly curved CCTs in heterogenous tissues (**Extended Data Fig. 7**). The above data evidenced that native single cytocapsular oncocells can generate super-long CCTs with variable superstructures, and that CCTs elongate along the path with least resistance in heterogeneous tissues *in vivo* (**Figs. 1 and Extended Data Fig. 5-7**).

In most instances, in invasive breast carcinoma tissues, there are large quantities of sectioned, short and curved CCT fragments. The high percentages of short CCT fragments with curves/turns/coils/ spirals suggest the high frequency of CCT elongation direction changes (**Fig. 1C-D**). The CCT elongation direction changes at CCT turns, curves, or crests/troughs of CCT wave fragments are caused by heterogeneous obstacles from tissue cells, ECMs, microarchitectures, and extend along the path with pores, microenvironment space, less cell density, less ECM density, less CCT density, where have least physical resistances (**Fig. 1C-D**).

In summary, the above *in vitro* and *in vivo* data on CCT elongation demonstrated that CCT elongation procedure/cycle contains several successive steps: 1) generation of enlarged NP via elongation and enlargement of nanoprotrusions (NPs) with least resistance, 2) enlarged NP grows into initial cytocapsular tubes (ICCTs), 3) with cytocapsulasome fusion, ICCT increase cytocapsular membrane areas, oncocell push and deform CCT membranes, and elongate CCT (**Extended Data Fig. 8**).

Consequently, CCT elongation, cancer cell metastasis via CCTs, and CCT degradation in tissues usually occur ahead of CCT invasion into blood vessels, as blood vessel walls have tight and dense cell-cell interactions and strong physical resistances (**Extended Data Fig. 9**).

### CCT branching morphogenesis increases CCT number and drives cancer metastasis

In some instances, at 72h after Bxpc3 pancreas cancer cell implantation, nanoprotrusions (NPs) at the side of CCTs enlarge and grow into branch enlarged NPs (branch ENPs, **Fig.S10A-B**). Subsequently, the branch ENPs develop into a branch initial CCTs (branch ICCTs) and branch CCTs (**Extended Data Fig. 10A-B**). In the examined 75 cytocapsular oncocell CCT with branches *in vitro* (21 of Bxpc3 pancreas cytocapsular oncocells, 18 of A549 lung cytocapsular oncocells, 36 of MDA-MB-231 breast cytocapsular oncocells), all these CCT branches employ bifid branching format.

Consistently, in clinical cancer tissues, CCT branching morphogenesis utilizes CCT side NP enlargement and bifid branching format (**Fig. 2A**). In the examined 126 CCTs with branch CCTs (55 CCT branches from breast carcinoma, 26 CCT branches from colon carcinoma, 17 CCT branches from prostate carcinoma, 28 CCT branches from lung carcinoma) employ side NP enlargement and bifid branching format. One stem CCT can perform multiple branching in a short CCT distance (**Fig.2A4**). After CCT remodeling, the branch CCT sizes in diameter/width are almost similar as that of stem CCTs. Oncocells bi-directionally migrate in CCT networks containing stem and branch CCTs (**Figs. 2A-B, and Extended Data Fig. 10C**).

**Fig. 2.**
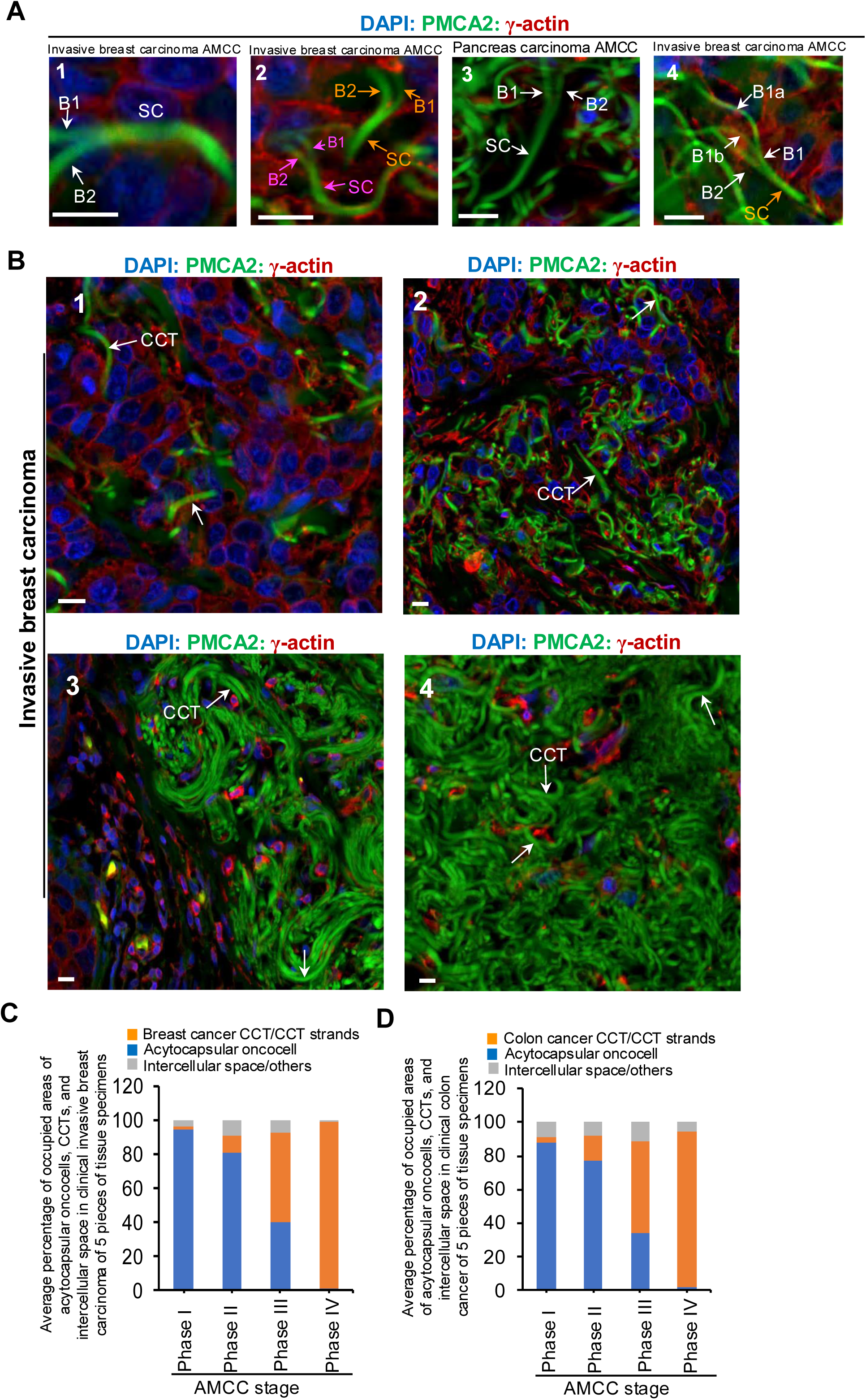
Cytocapsular tube bifid branching morphogenesis significantly increase CCT number and promote CCT conducted cancer metastasis. (**A**) Representative IHC fluorescence microscope images of bifid CCT branching morphogenesis format in breast and pancreas cancer tissues *in vivo*. Panel A1, a breast cancer stem CCT (SC) with two branch CCTs (branch 1 (B1) and branch 2 (B2)). Panel A2, two breast cancer stem CCTs (CSs) and each CS with two branch CCTs. Panel 3, a pancreas stem CCT with two branch CCTs. Panel 4, a breast cancer stem CCT (SC) with two branch CCTs (B1 and B2), and the short branch 1 (B1) continuously develops into two CCT branches (B1a and B1b). (**B**) Representative IHC fluorescence microscope images of invasive breast carcinoma in 4 AMCC phases: phase I, II, III and IV: panel B1 (AMCC phase I), panel B2 (AMCC phase II), panel B3 (AMCC phase III), and panel B4 (AMCC phase IV). CCT branching morphogenesis significantly increase CCT numbers, and promote acytocapsular oncocells invade into CCTs via alloentry followed by CCT-conducted cancer ell metastasis. Scale bar, 10μm. (**C**) Quantitative statistical analysis of occupied areas of breast cancer CCT/CCT strands, acytocapsular oncocells, intercellular spaces/others (such as micro blood vessels) in the 4 AMCC phases. (**D**) Quantitative statistical analysis of occupied areas of colon cancer CCT/CCT strands, acytocapsular oncocells, intercellular spaces/others (such as micro blood vessels) in the 4 AMCC phases.

In AMCC stage in clinical breast and colon carcinoma tissues, CCT branching morphogenesis significantly increase CCT numbers from AMCC phase I to AMCC phase IV (**Fig.2**). CCT branching morphogenesis contributes significant increase in both CCT number and density in AMCC stage in breast cancer, prostate cancer and colon cancer (**Extended Data Fig. 11**). CCT branching morphogenesis is universally presented in all the checked 288 subtypes of cancers in 35 kinds of human tissues and organs (**Extended Data Fig. 4**). In addition, increased CCT occupied areas associate with increased cancer cell metastasis, and decreased left cancer cells (**Fig. 2C-D, and Extended Data Fig. 11**). These data demonstrated that CCT branching morphogenesis substantially contributes to CCT number and density increase, cancer cell metastasis progression, and cancer evolution.

### CCT regeneration and elongation drive continuous cancer metastasis in AMCC

In acytocapsular oncocell mass-CC/CCT complex (AMCC) stage, acytocapsular oncocell continuously perform uncontrolled oncocell proliferation, and one generation of CCTs can only reach metastasis of parts of acytocapsular oncocells, and massive cancer cells remain (**Figs. 3 and Extended Data Fig. 12**). Interestingly, after old CCTs degrade, decompose, and disappear, leaving CCT cavity strips formed 3D CCT cavity networks in acytocapsular oncocell mass (**Figs.3 and Extended Data Fig. 12**). Subsequently, new CCTs are regenerated. With CCT branching morphogenesis, CCT number and density significantly increase, and extend along CCT cavity strips formed 3D CCT cavity networks left by previous generation CCT network (**Figs.3 and Extended Data Fig. 12**). The new generation CCT networks continuously let more oncocells dissemination and leave away, and therefore drive cancer relapse (**Figs.3 and Extended Data Fig. 12**). The phenomena of repeated CCT regeneration and elongation universally present in all checked 289 subtypes of cancers in 34 kinds of human tissues and organs (**Extended Data Fig. 4**). These observations demonstrated that repeated CCT regeneration and elongation in AMCC drive repeated and continuous cancer metastasis.

**Fig. 3.**
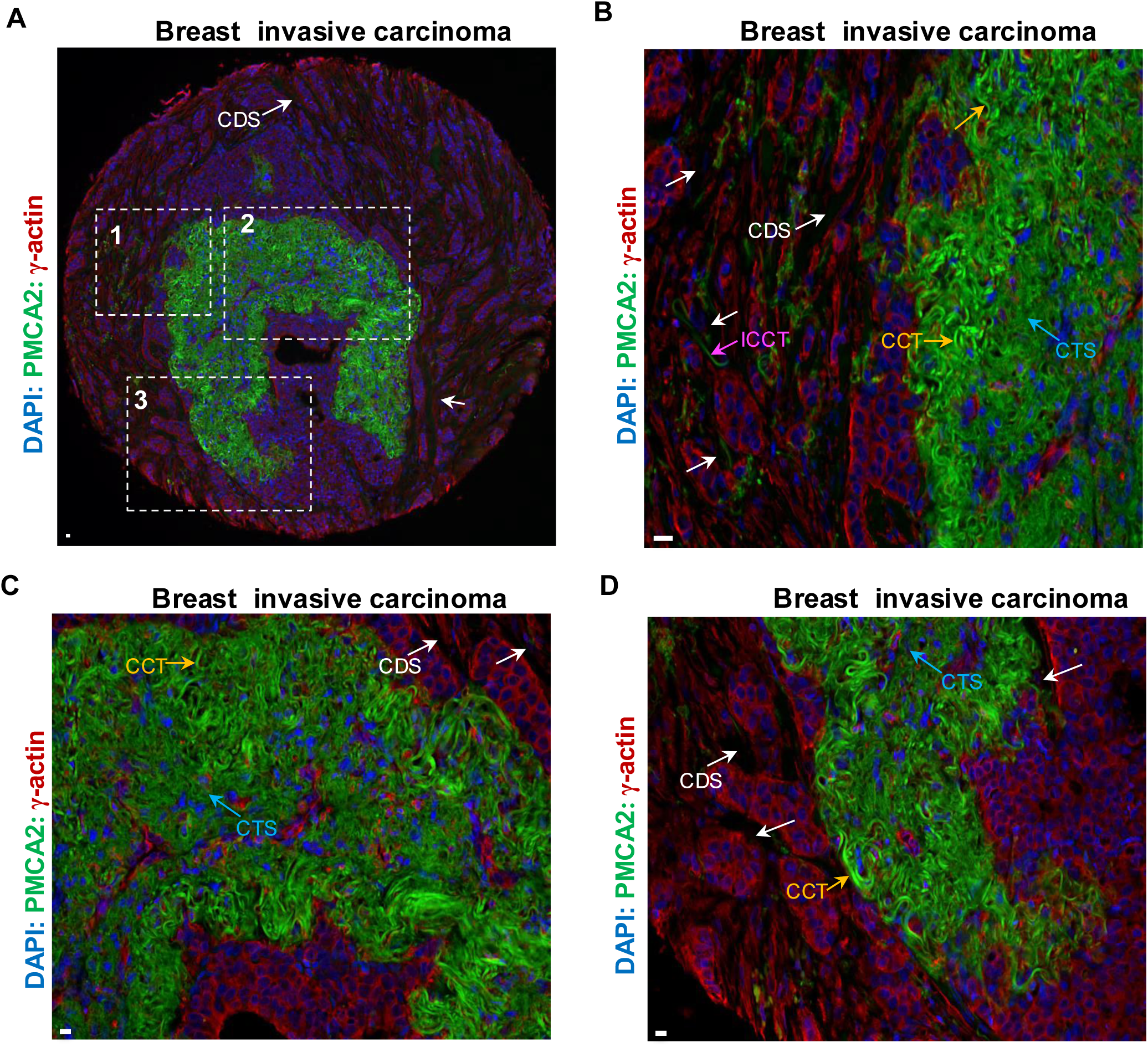
CCT regeneration in AMCC promotes new cycles of CCT-conducted cancer metastasis. (**A**) Representative breast invasive carcinoma in AMCC stage with CCT Degradation left cavity Strip (CDS, white arrows) interconnection formed CDS networks. There are many CCT degradation left cavity strips (CDSs) in variable widths crossing the dense acytocapsular oncocell masses in AMCC, and separate dense acytocapsular oncocell (AO) masses into many dense AO mass islands. The CCTs degrade into cloud-like CCT strands (smear green cloud in the image) followed by disappearance. The framed areas (1, 2, and 3) are enlarged and shown in the panels B, C, and D, respectively. (**B**) Enlarged framed area 1 in (A). Many newly regenerated ICCTs (purple arrows) and CCTs (orange arrows) elongate along and fill the previous CDSs (white arrows). (**C**) Enlarged area 2 in (A). Massive newly regenerated CCTs fill CDSs (white arrows) and many cytocapsular oncocells invade into regenerated CCTs and leave aways. There are some regenerated CCTs in degradation into CCT strands (CTSs, cyan arrows). (**D**) Enlarged area 3 in (A). Massive newly regenerated CCTs fill CDSs, in which regenerated CCTs tightly contact with dense AO islands, and allow AOs invade into CCTs via alloentry, followed by continuous cancer metastasis in AMCC. Scale bar, 10μm.

### CCT superstructures promote complete cancer metastasis in AMCC stage

Single, super-long, and highly curved/coiled CCTs generate big, triangle-like CCT super-structures (**Fig.4**). Multiple big, such CCT super-structures compose multi-layered CCT super-structure complex, wrapping multiple irregular shaped cytocapsular tumors in variable sizes (**Fig.4**). Similar multiple-layered CCT superstructure complexes in diverse morphologies are present in pancreas carcinoma, lung carcinoma, prostate carcinoma (**Extended Data Fig. 13**). The acytocapsular oncocell densities in the diverse mega breast CCT super-structures are close to or already zero, (**Fig.5**), suggesting that mega CCT superstructure complexes significantly enhance CCT density, dramatically increase CCT membrane surface areas for acytocapsular oncocell alloentry into CCTs, and therefore promote cancer metastasis via CCTs. Mega CCT superstructure complexes allow acytocapsular oncocell masses to completely leave away via CCT conducted cancer disseminate (**Fig.5**). Therefore, CCT superstructure complexes advance in reaching complete cancer metastasis of acytocapsular oncocell masses.

**Fig. 4.**
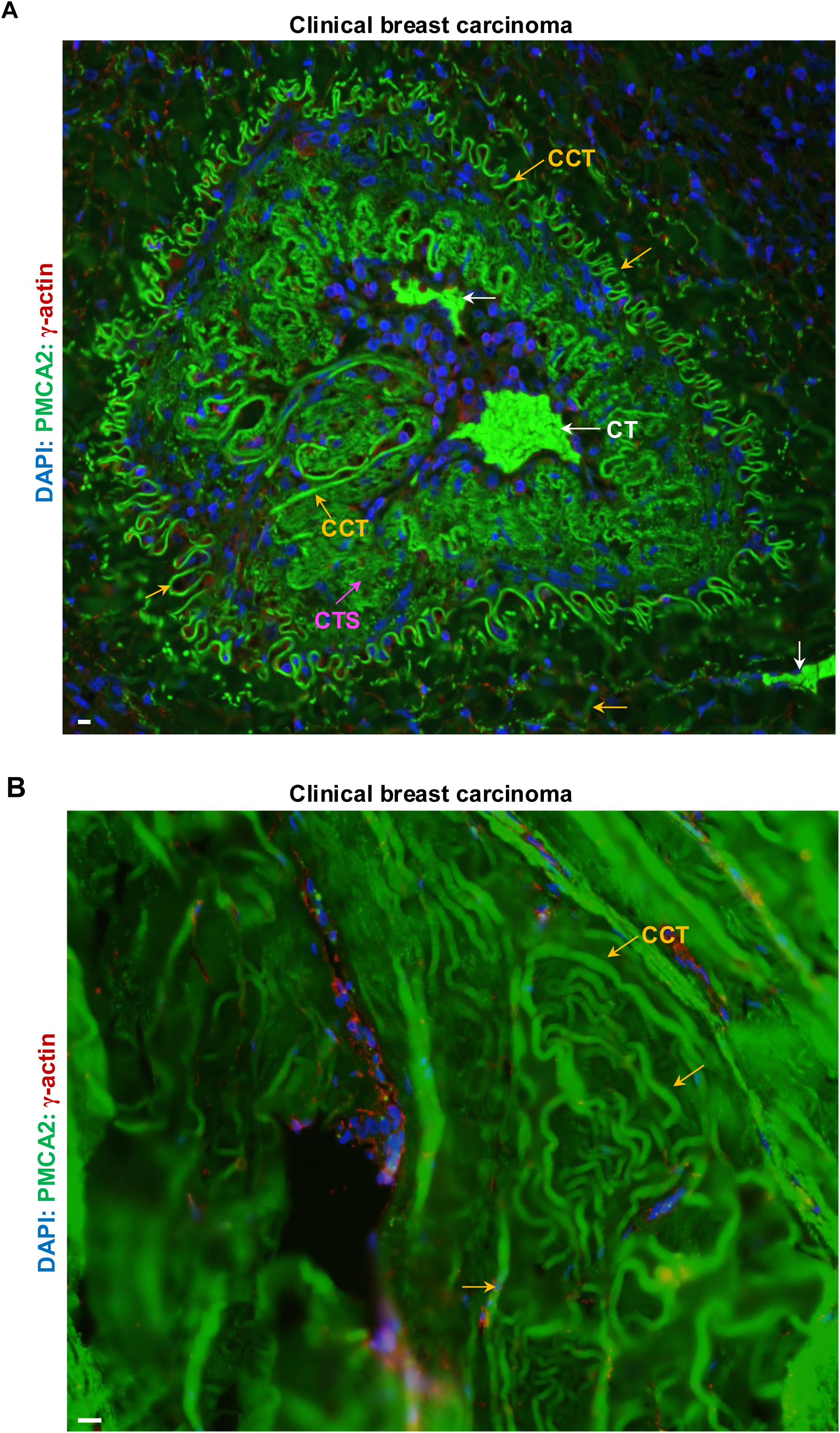
CCT mega superstructures promote cancer metastasis *in vivo*. (**A**) Representative IHC fluorescence image of a super-large breast CCT mega superstructure complex with two-layered, irregular triangle-shaped, and highly curved CCT superstructures with many and dense CCT curves/coils of each CCT superstructure. There are two irregular-shaped cytocapsular tumors (CTs, white arrows) in the CCT mega superstructure. Many acytocapsular oncocells (AOs) are left via mega CCT superstructure conducted metastasis. Some CCTs (orange arrows) degrade into CCT strands (CTSs, purple arrows). (**B**) Representative image of super-large, many-layered, highly dense CCT formed mega CCT superstructures in the late breast carcinoma AMCC stage. Most acytocapsular oncocells (AOs) are left via mega CCT (orange arrows) superstructure conducted metastasis. CCT mega superstructures promote complete cancer metastasis of AMCCs. Scale bar, 10μm.

**Fig. 5.**
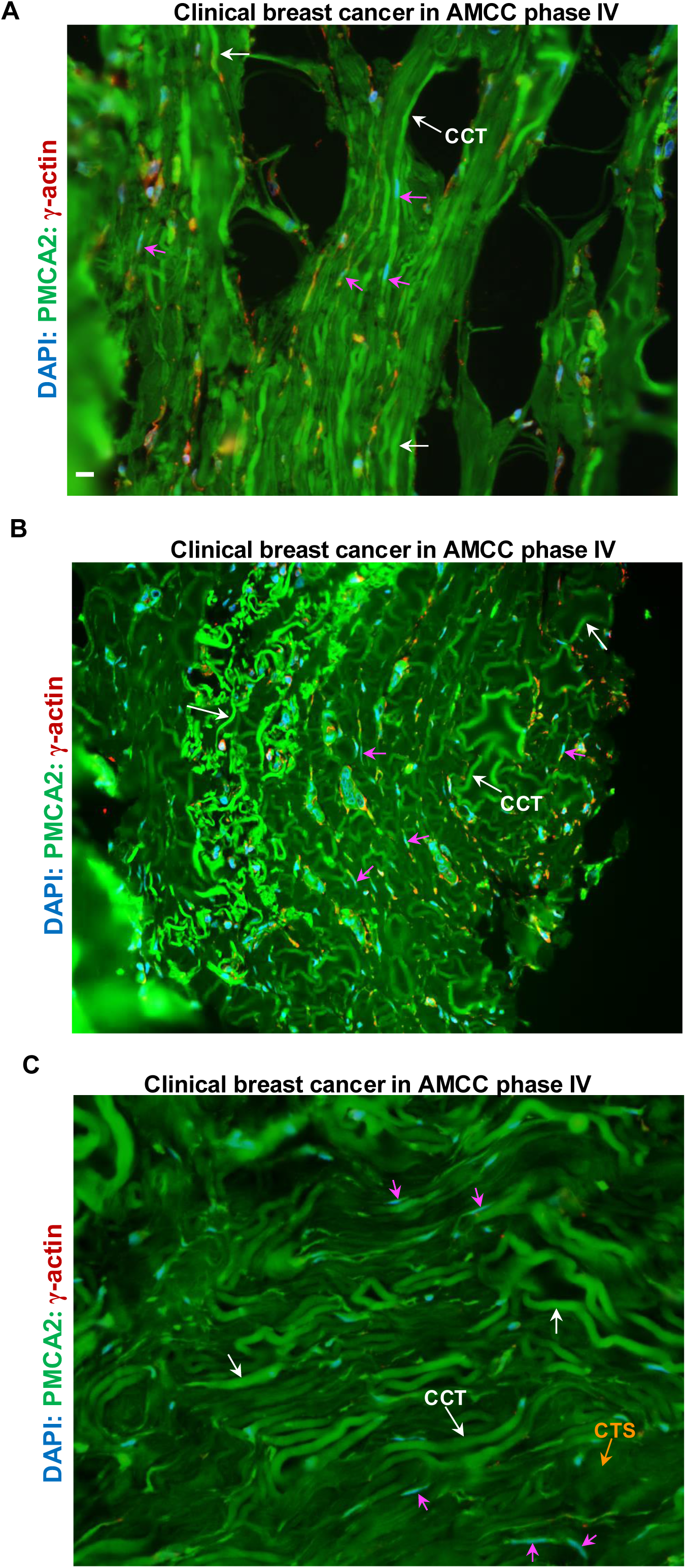
Diverse CCT mega superstructures conduct complete cancer metastasis in AMCC stage *in vivo*. (**A**) Representative image of super-large and dense CCT bunch interconnection formed mega CCT network superstructures in the breast cancer AMCC phase IV. All acytocapsular oncocells (AOs) are left away via mega CCT superstructure conducted metastasis. (B) Representative image of super-large, many-layered, highly curved/coiled CCT formed mega CCT superstructures in the breast cancer AMCC phase IV. All acytocapsular oncocells (AOs) are left away via mega CCT superstructure conducted metastasis. (**C**) Representative image of super-large, many-layered, highly dense CCT formed mega CCT superstructures in the breast carcinoma AMCC phase IV. All acytocapsular oncocells (AOs) are left via mega CCT superstructure conducted metastasis. Some CCTs degrade into CCT strands (CTSs, orange arrows). CCTs (white arrows) and intracytocapsular oncocells in migration in CCTs (purple arrows, sectioned and nuclei are exposed) are shown. Scale bar, 10μm.

## Discussion

The discovery of cytocapsula, cytocapsular tube (CCT), CCT networks, cytocapsular oncocells/tumors, and cytocapsular tumor network systems (CTNSs) in our two initial reports on CCT-cancer have unveiled a previously unnoticed principle of native cancer *in vivo*. Here, we show that CCT elongation in the least resistance path and CCT branching morphogenesis in bifid format universally present in cancers *in vivo*. This was achieved via a 7-year effort to collect >300,000 pieces of CCT fluorescence microscope images of clinical cancer tissue specimens from >15,000 cancer patients from tissue banks worldwide, with a goal to investigate CCT development in 288 subtypes of human cancers in diverse stages. The CCT-elongation mechanisms that emerged as principles for native cancer progression research, and clinical cancer diagnosis and therapies.

The advantages of CCT elongation via nanoprotrusion (NP) enlargement conducted new CCT formation are: 1) the single NPs are in tiny sizes of 21nm ∼ 82nm in diameter/width, much smaller than CCTs in sizes (3-10 μm in diameter/width), which help NPs to extend into large quantities of micropores (<100nm in width) in microenvironments with minimum resistance; 2) the large quantities of NPs surrounding CCTs and CCT leads help CCTs to sense resistance in all 3D directions in micro positions in local microenvironments, and choose the NPs with least resistances to enlarge and develop into new CCTs; 3) NP membranes are extended from CCT membranes and share similar cytocapsular membrane structural components (membrane lipids, membrane proteins, beneath microfilament networks, and so on), and relatively easy to enlarge in sizes; 4) the open connection of NPs at/in CCT ultrastructural complexes facilitate NPs to access and obtain intracytocapsular oncocell released membrane building materials and proteases, without physical barriers or obstacles; 5) NPs in the leading release proteases and proteolyze to remove resistances from ECMs (including bone matrices), cell-cell interactions (even brain-blood barriers) and cell-ECM interactions in tiny scale initially, which is an efficient way to find out a path with least resistance. The above multiple unique advantages significantly facilitate NP-conducted CCT elongation in the least resistance path to overcome/remove heterogeneous macro and micro resistances in heterogeneous human tissues/organs

The CCT-elongation mechanisms elucidate the principles of CCT elongation, CCT network expansion, CCT freeway system conducted and cytocapsular membrane protected cancer metastasis, and intro-cytocapsular oncocell proliferation and migration driven cancer progression in heterogeneous human tissues and organs. The leading intro-cytocapsular oncocells drive CCT elongation and CCT network expansion in multiple aspects: 1) generate and release cytocapsulasomes and other membrane-enclosed tiny bodies as CCT elongation membrane building blocks; 2) release proteases and provide proteolysis to remove resistances from ECMs (including bone matrices), cell-cell interactions (including brain-blood barriers) and cell-ECM interactions; 3) receive signals from outside nanoprotrusions and decide the preferable NPs with least resistance to enlarge and grow up; 4) generate blebs to sense extracytocapsular microenvironmental resistances and push neighboring cells/ECMs back to make space for NP enlargement and initial CCT expansion in diameter/width and length; 5) push and deform cytocapsular membranes into tube-shaped morphologies, and generate new NPs anchoring CCT in place; 6) drive side NPs to develop into branch CCTs and promote CCT branching morphogenesis, and therefore provide expanded 3D CCT freeway systems for oncocell metastasis in a membrane barrier protected cell dissemination format, escape from immune cell attack, achieve pan-cancer drug resistance; 7) Drive repeated CCT regeneration, and new generations of CCTs and CCT networks promote new cycle of cancer metastasis, resulted in repeated tumor relapse.

CCT branching morphogenesis in side NP enlargement and bifid format is propertied with multiple characterizations: 1) branch CCTs are developed from CCT side and enlarged NPs with least resistance, which interconnect with stem CCTs with open connection, sharing with similar cytocapsular membrane components and shared the similar/same materials in stem CCT lumens and NP lumens, 2) intracytocapsular oncocells (in long, thin and spindle-shaped mesenchymal morphologies in initial CCTs) release cytocapsulasomes, which integrate into initial CCTs, and increase CCT membrane areas; push and deform cytocapsular membranes to form bigger tube-shaped CCTs; 3) stem CCT sides have large quantities of CCTs already extend into ECMs anchoring CCTs in place, which provide substantial opportunities to develop multiple or many CCT branches via multiple/many times of bifid branching; 4) CCT branching morphogenesis significantly increase CCT numbers and expand CCT networks in network sizes and cytocapsular membrane area sizes for outside acytocapsular oncocells to invade into CCT network systems via alloentry format. Therefore, more CCTs promote cancer cell metastasis to leave away from the stressful inner tumor microenvironments (uncontrolled cancer cell proliferation caused nutrient deprivation, hypoxia, low *p*H, elevated metabolism waste molecules, insufficient growth factors, high competition from neighboring cancer cells, toxic microenvironments) for better survival conditions.

In summary, the CCT elongation laws and mechanisms of CCT elongation and CCT branching morphogenesis described here may facilitate native cancer research in a CTNS level and help clinical cancer applications, including cancer diagnosis, surgery, caner drug research and development, and CTNS targeted effective cancer therapy.

## Acknowledgements and funding sources

This work was funded by: Cytocapsula Research Institute Fund (T. Y.) and Centiver Ltd Fund (T. Y. and G.W.). This work was supported by Cellmig Biolabs Inc. All data produced in the present work are contained in the manuscript. We thank Centiver Ltd. team and Cellmig Biolabs Inc. team for the help on CCT analyses. We greatly acknowledge Dr. Ed Harlow of Harvard Medical School (USA) and Cancer Institute of University of Cambridge (UK), Dr. Nahum Sonenberg of McGill University of Canada, Dr. Michael Roehrl of Beth Israel Deaconess Medical Center (BIDMC), Dr. Stephen Ganshirt of Northwest University Medicine Hospital, Dr. Joan Brugge of Harvard Medical School, for their help and meaningful discussion in the study.

## Author contributions

T.Y. designed research; T.Y. Resources; T.Y. performed research; T.Y. supervision and project administration; T.Y. and G.W. analyzed data; T.Y. and G. W. wrote the manuscript.

## Conflict of interest statement

The authors are cofounders of and hold equity in Centiver Ltd and Cellmig Biolabs Inc. to support future basic research on cytocapsular oncocells and cytocapsular tumor network systems.

## Extended Data

### Glossary of abbreviations and acronyms

CC: Cytocapsula
CCT: Cytocapsular Tube
CT: cytocapsular tumor/tumorsphere
PCT: Prophase Cytocapsular Tumor
NT: Nuperphase (late) Tumor
AMCC: Acytocapsular Oncocell Mass-CT/CCT Complex
NP: Nano Protrusions

## Materials and Methods

### Reagent, Antibodies, Clinical Cancer Tissues, and Devices

CC/CCT culture kits (Celldevi, https://www.celldevi.com, Cat. CD0104, six-well plates; Cat. CD0105, 12-well plates, and Cat. CD 0106, 24-well plates) and kits with glass cover slips in the well bottom and embedded by CC/CCT culture matrix layer (Cat. CD 0112, six-well plate) were ordered from Celldevi Inc. CC/CCT culture kit fixation kit (Cat. CD0201) were ordered from Celldevi Inc. CT-M2 fluorescence microscope (with high resolution, high signal/noise ratio, low background noise CCT imaging) for CCT fluorescence microscope imaging and CCT analysis software were ordered from Celldevi. Cancer cell lines of pancreas cancer cell Bxpc3, breast cancer cell MCF-7 and colon cancer cell SK-CO-1 and cell culture media were ordered from ATCC. Breast cancer stem cells (BCSCs) of HMLER(CD24^low^/CD44^high^)^FA^ subpopulation cells were self-isolated by flow cytometry and cultured. Rabbit anti-PMCA2 antibody (polyclonal, ab3529; 1:200 dilution), mouse anti-γ-actin antibody (monoclonal, ab123034; 1:200 dilution) for immunofluorescence assay were ordered from Abcam. DAPI (1:1,000 dilution in the immunofluorescence assay) was ordered from KPL. Human normal and cancer tissue specimens described here were ordered from Tissue Array and US Biolabs and were deidentified prior to use. JEOL 1200EX Transmission electron microscope or a Tecnai G2 Spirit BioTWIN microscope, and images were recorded with an AMT 2k CCD camera.

### Cytocapsular tumorsphere and cytocapsula growth *in vitro*, immunohistochemistry staining and imaging

Pancreas cancer Bxpc3 cells were implanted in CC/CCT culture kit (Cat. CD 0112, Celldevi) following the kit manual. At 36h, Bxpc3 cells generated cytocapsulas (CCs). Some cytocapsular oncocells performed ecellulation. Cytocapsular oncocells and ecellulated CCs were performed fixation kit and immunohistochemistry staining. At different time of 48h, 72h, 68h, 74h, 78h, 84h, 96h, 108h after cell implantation, Bxpc3 cancer cells engender cytocapsular oncocells, and grow into cytocapsular tumorspheres in different sizes with CC tightly wrapping oncocell mass or with wide cytocapsular lumens, and ecellulation of cytocapsular tumorspheres. These cytocapsular tumorspheres and ecellulated cytocapsular tumorspheres were fixed by Celldevi Inc. CC/CCT fixation kit (Celldevi, CD0201) in the 6-well plate, and then taken out and put onto slides, followed by immunohistochemistry (IHC) staining.

### Cytocapsular tumorsphere and cytocapsula growth *in vitro*, immunohistochemistry staining and imaging

IHC staining was performed with rabbit anti-PMCA2 polyclonal primary antibodies (1:200 dilution), mouse anti-γ-actin monoclonal primary antibodies (1:200 dilution), Goat anti-Mouse IgG (H+L) Highly Cross-Adsorbed Secondary Antibody, Alexa Fluor Plus 555 (Thermo Fisher), and Goat anti-Rabbit IgG (H+L) Highly Cross-Adsorbed Secondary Antibody, Alexa Fluor Plus 488, Thermo Fisher), and DAPI staining (1:000 dilution). Fluorescence images were taken with a Nikon 80i upright microscope with a 20× or 40× lens. All images were obtained using MetaMorph image acquisition software and were analyzed with ImageJ software.

### CCT Histology and Immunohistochemical Staining Analysis

The 9972 formalin-fixed, paraffin-embedded (FFPE) human cancer tissue specimens (4-5μm in thickness) from 9784 cancer patients, 14 human normal tissue FFPE specimens form 14 patients, and 126 human benign tumor tissue FFPE specimens from 126 patients were processed immunohistochemistry and hematoxylin and eosin (H&E) staining. Immunohistochemical fluorescence tests were performed to stain cytocapsular tubes using rabbit anti-PMCA2 polyclonal primary antibodies (1:200 dilution), mouse anti-γ-actin monoclonal primary antibodies (1:200 dilution), Goat anti-Mouse IgG (H+L) Highly Cross-Adsorbed Secondary Antibody, Alexa Fluor Plus 555 (Thermo Fisher), and Goat anti-Rabbit IgG (H+L) Highly Cross-Adsorbed Secondary Antibody, Alexa Fluor Plus 488, Thermo Fisher), and DAPI staining (1:000 dilution). Fluorescence images were taken with a Nikon 80i upright microscope with a 20× or 40× lens. All images were obtained using MetaMorph image acquisition software and were analyzed with ImageJ software.

### Imaging Acquisition

DIC and fluorescence images of fixed cells (with or without cytocapsulae) were taken with an 80i upright microscope and a digital Hamamatsu ORCA-ER cooled CCD camera with a 20× or 40× lens. The bright-field phase-contrast image was taken using a Nikon digital camera. The cytocapsula initiation ratio per high-performance field (HPF; 200×) and the number of elongated cytocapsulae per high-performance field were quantified. All images were obtained using MetaMorph image acquisition software and were analyzed with ImageJ software.

### Transmission Electron Microscope

The 6-well plate CC/CCT culture kits were ordered from Celldevi. Breast cancer stem cells (BCSCs) of HMLER(CD24^low^/CD44^high^)^FA^ subpopulation cells were self-isolated by flow cytometry and cultured. BCSC cultures with cytocapsulas and nanoprotrusions (NPs) in the kits (>40 μm in depth) were fixed with 1:1 mixtures of formaldehyde-glutaraldehyde-picric acid fixative (2.5% paraformaldehyde, 5.0% glutaraldehyde, 0.06% picric acid in 0.2 M cacodylate buffer):cell culture medium. The cells and cytocapsulae and NPs were then postfixed for 30 min in 1% osmium tetroxide (OsO4)/1.5% potassium ferrocyanide (KFeCN6), washed three times in water, and incubated in 1% aqueous uranyl acetate for 30 min followed by two washes in water and subsequent dehydration in grades of alcohol (50, 70, 95, and 2 × 100%; 5 min each) (1). Cells and cytocapsulae with NPs were infiltrated for 2 h to overnight in a 1:1 mixture of propylene oxide and TAAB Epon (Marivac Canada Inc.). The samples were subsequently embedded in TAAB Epon and polymerized at 60 °C for 48 h. Ultrathin sections (∼60 nm in thickness, no curved or overlapping sections) were cut on a Reichert Ultracut S microtome, picked up on copper grids, stained with lead citrate, and examined in a JEOL 1200EX Transmission electron microscope or a Tecnai G2 Spirit BioTWIN microscope, and images were recorded with an AMT 2k CCD camera.

### Nanoprotrusion bunches in bright filed and fluorescence microscope images *in vitro*

Nanoprotrusions (NPs, 21nm ∼ 82nm in diameter/width) of CCs/CCTs are in visible in bright field and fluorescence microscope due to the limited resolution of these microscopes. The NP bunches with big sizes (100nm ∼ 152nm in width) are visible in high resolution bright filed microscope images. The NP bunches with big sizes (100nm ∼ 152nm in width) and coated with thin or no glycocalyx layers are visible in CT-M2 fluorescence microscope images with high resolution, high signal/noise ratio and low background (but invisible in common fluorescence microscope).

### Initial cytocapsular tube (ICCT) and CCT analysis in cancer tissues *in vivo*

In cancer tissues, initial cytocapsular tubes (ICCTs) are small (0.2μm ∼ 0.5μm in diameter/width) straight/curved cytocapsular tubes without degradation, with or without long and thin cancer cells in migration inside. Cytocapsular tubes (CCTs) are established cytocapsular membrane enclosed tubes (0.5μm ∼ 10μm in diameter/width). CCT degraded strands show phenomena of: networks of very thin strands, dot array collapse membrane degradation, cloud line-status.

### Data collection

Cytocapsular tubes (CCTs, not sectioned, longitudinally sectioned, and cross sectioned) without degradation (3∼6μm in measured diameter) were counted using a fluorescence microscope and ImageJ. The presence of CCTs degrading into thick strands (1∼2μm in measured diameter), thin strands (0.2∼1μm in measured diameter), or the disintegration state were reported without quantification. The patients providing formalin-fixed paraffin-embedded tissue samples gave informed consent that they understood that the biopsies (needle biopsy or surgical biopsy, or autopsy tissues from US Biomax) were performed for *in vitro* research purposes only. Comparative deidentified samples of normal tissues, benign tissues, carcinoma *in situ*, cancer, paracancer, metastatic tissues with their cancer stages identified according to the tumor (T), node (N), and metastasis (M) TNM system (cancer stages: 0, I, II, III, IV) were obtained from archival materials (Tables S1). The cancer, paracancer and metastatic cancer tissues, in which the original cancer niches were identified by indicated cancer specific molecular markers, were identified by hospital pathology laboratories and obtained from archival material. Autopsy tissues samples have many post-life CC/CCT degradation and CCs/CCTs will not be quantified. Biopsy samples from FFPE with fresh tissues present high fidelity of CC/CCT status and CCs/CCTs are quantified and reported.

### Quantification and Statistical Analysis

The statistical methods used for comparisons are indicated in the relevant Figure legends and in the sections below. The diameters, widths, and lengths of cytocapsulae and cytocapsular tubes were measured with MetaMorph or ImageJ. The diameters, widths, and lengths of sectioned nanoprotrusions and nanoprotrusion bunches were measured with transmission electron microscope and its software. For cytocapsular oncocell and cytocapsular tumorsphere assays, at least 20 cytocapsular oncocell or cytocapsular tumorsphere were measured per condition, and two-tailed Student’s test was used to determine statistical significance. The graph plots are mean ± SD. In initial CCT analysis in cancer types/subtypes, at least 3 samples per cancer subtype were checked. In cytocapsular tube quantitation assays, for each specimen, the number of fully intact cytocapsular tubes was counted in 5 areas (0.35mm x 0.35mm, length x width) of the sample (top, bottom, left, right, and center), and the cytocapsular tube density (CCT/mm^2^) was calculated and determined for each area. The average CCT density across the 5 sites was treated as the specimen’s overall CCT density and round up to digits. The checked tissues sample numbers of CCTs with turns/waves/curves, CCT/CCT strands occupied area percentages, CCT regeneration in AMCC stages are shown in the text (or figure legends).

## Extended Data Figures and Legends

**Extended data Fig.1.**
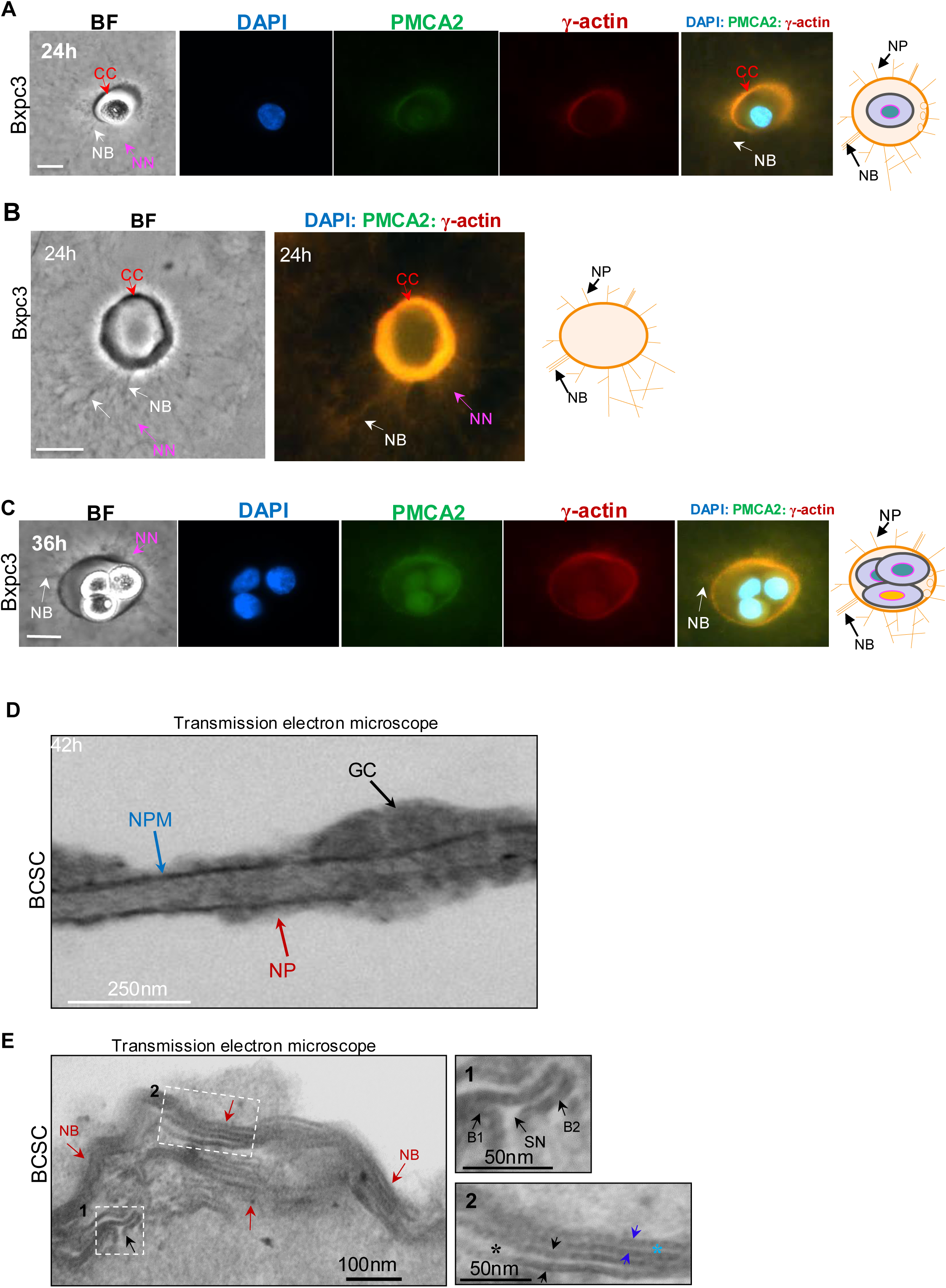
Cytocapsular oncocells and tumorspheres generate cytocapsular nanoprotrusions (NPs), and ultrastructure of cytocapsular nanoprotrusions (NPs) and NP bunches *in vitro*. **(A)** Representative bright field (BF) and immunohistochemistry (IHC) fluorescence microscope images of a single cytocapsular oncocell (CO) with cytocapsular nanoprotrusions (NPs) at 24h after Bxpc3 pancreas cancer cell implantation in 3D CC/CCT culture kits. A schematic diagram of single COs with NPs and NP bunches. **(B)** Representative bright field and IHC fluorescence microscope images of ecellulated cytocapsulas (ECs) with NPs and NP bunches of single Bxpc3 cytocapsular oncocells in 3D CC/CCT culture kits. The ECs collapse, and form deflated, concave discs. NP networks (NNs) interconnect with and surround the acellular cytocapsulas. A schematic diagram of single ecellulated cytocapsula with NPs and NP bunches. **(C)** Representative bright field and IHC fluorescence microscope image of a Bxpc3 pancreas cancer cytocapsula tumorsphere with NPs and CNP bunches. A schematic diagram of single cytocapsular tumorsphere with CNPs and CNP bunches. The cytocapsula (CC, orange arrows), cytocapsular nanoprotrusion (NP), cytocapsular nanoprotrusion bunches (NB, white arrows), cytocapsular nanoprotrusion network (NN, purple arrows), incytocapsular oncocell (ICO, purple arrows) are shown. Scale bar, 10μm. **(D)** Representative transmission electron microscope (TEM) image of longitudinal section of a single cytocapsular nanoprotrusion (NP) fragment at 42h after breast cancer stem cell (BCSC) HMLER(CD24^low^/CD44^high^)^FA^ subpopulation cell implantation in 3D CC/CCT culture kits. The NPs are coated with glycocalyx (GC, black arrows) layers in variable thickness. Cytocapsular nanoprotrusion (NP, red arrows), cytocapsular nanoprotrusion membrane (NPM, blue arrows) are shown. Scale bar, 250nm. **(E)** Representative TEM image of longitudinal section of cytocapsular nanoprotrusion bunches (NBs, red arrows). Multiple NPs align together tightly or loosely and form NBs in curved morphologies in thick glycocalyx layers. Scale bar, 100nm. The enlarged NBs are show in the right two panels: Panel E1, longitudinal section of a stem NP (SN) with two branch (B1 and B2) NPs in a bifid NP branching format with an open interconnection between the stem NP and the two NP branches; Panel E2, longitudinal sections of two NPs align together closely with the 4 sectioned and exposed NP membranes, which are embedded in glycocalyx layer. Asterisks in cyan and black colors: longitudinal sectioned lumens of the two NP lumens. Scale bar, 50nm.

**Extended data Fig.2.**
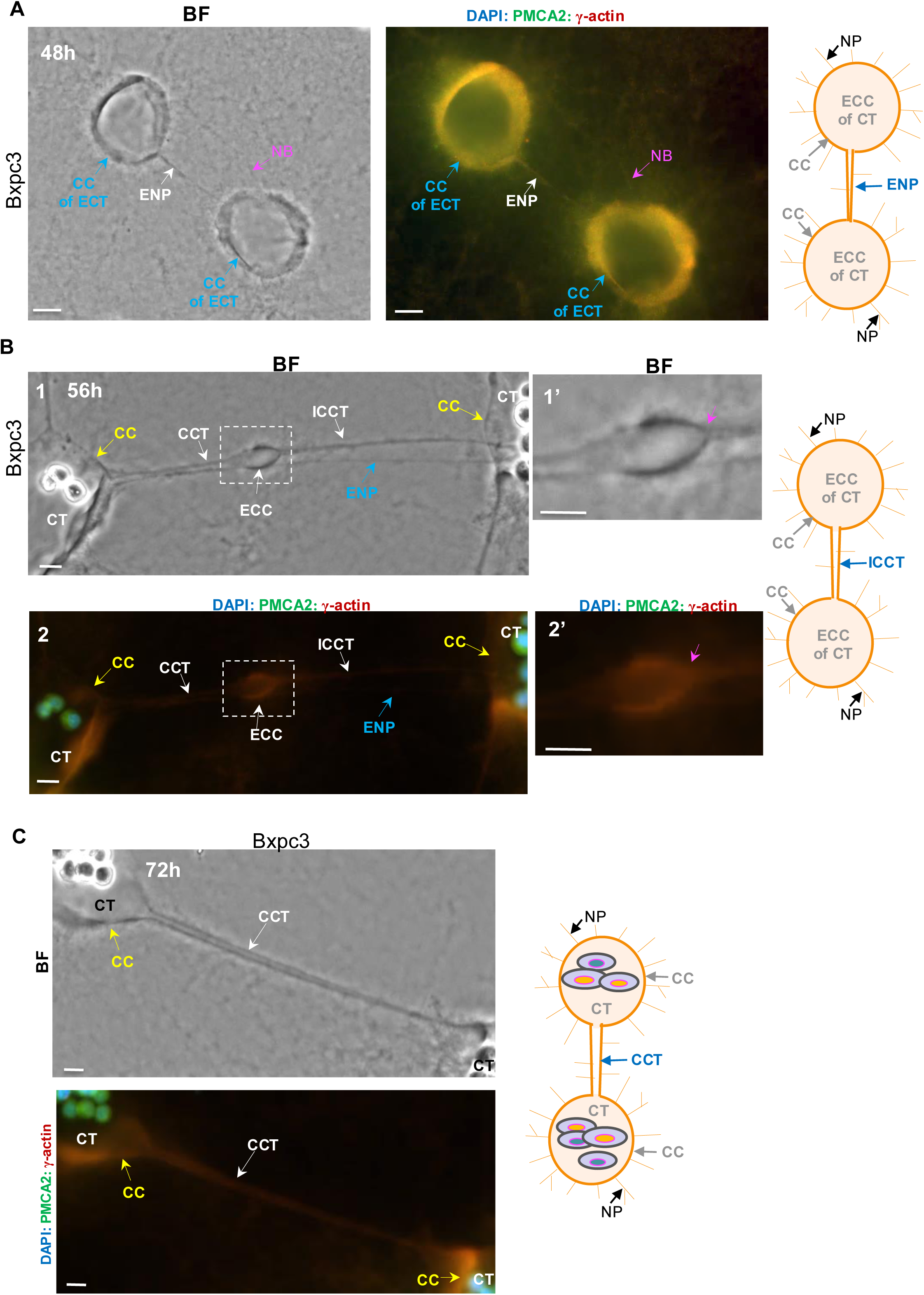
CCT nanoprotrusions (NPs) develop into enlarged NPs, followed by development into initial CCTs *in vitro*. (**A**) Representative bright field and IHC fluorescence microscope images show that cytocapsular nanoprotrusions (NPs) with shortest distance and least resistance between ecellulated cytocapsulas (ECCs) of cytocapsular tumorspheres (CTs) enlarge, and grow into enlarged cytocapsular nanoprotrusions (ECNs, white arrows). NP bunches (NBs, purple arrows) and cytocapsulas (CCs) of ecellulated CTs (ECTs, cyan arrows) are shown. (**B**) Representative bright field (BF) and IHC fluorescence microscope images show that in initial cytocapsular tubes (ICCTs) linking two cytocapsular tumorspheres (CTs), single Bxpc3 intracytocapsular oncocells migrate inside of ICCT, and make enlarged CCT bulge fragments behind. A single intracytocapsular oncocell ecellulated, and left an enlarged and oval-shaped CCT membrane bulge. The enlarged CCT membrane bulge is enlarged in panel B1’ and B2’. Purple arrows show the open connection between the CCT bulge front and ICCT. Enlarged NP (ENP, cyan arrow), cytocapsular tumorspheres (CTs) with the major intracytocapsular oncocells are ecellulated and only a few intracytocapsular oncocells left inside, and cytocapsula (CC, yellow arrows) of CT are shown. (**C**) Representative bright field (BF) and IHC fluorescence microscope images show that a long, straight CCT (white arrow) interconnects with two cytocapsular tumorspheres (CTs) with shortest distance. There are many NPs surrounding the CCTs. Cytocapsular tumorspheres (CTs) with the major intracytocapsular oncocells are ecellulated and only a few intracytocapsular oncocells left inside, and cytocapsula (CC, yellow arrows) of CT are shown. Scale bar, 10μm.

**Extended data Fig.3.**
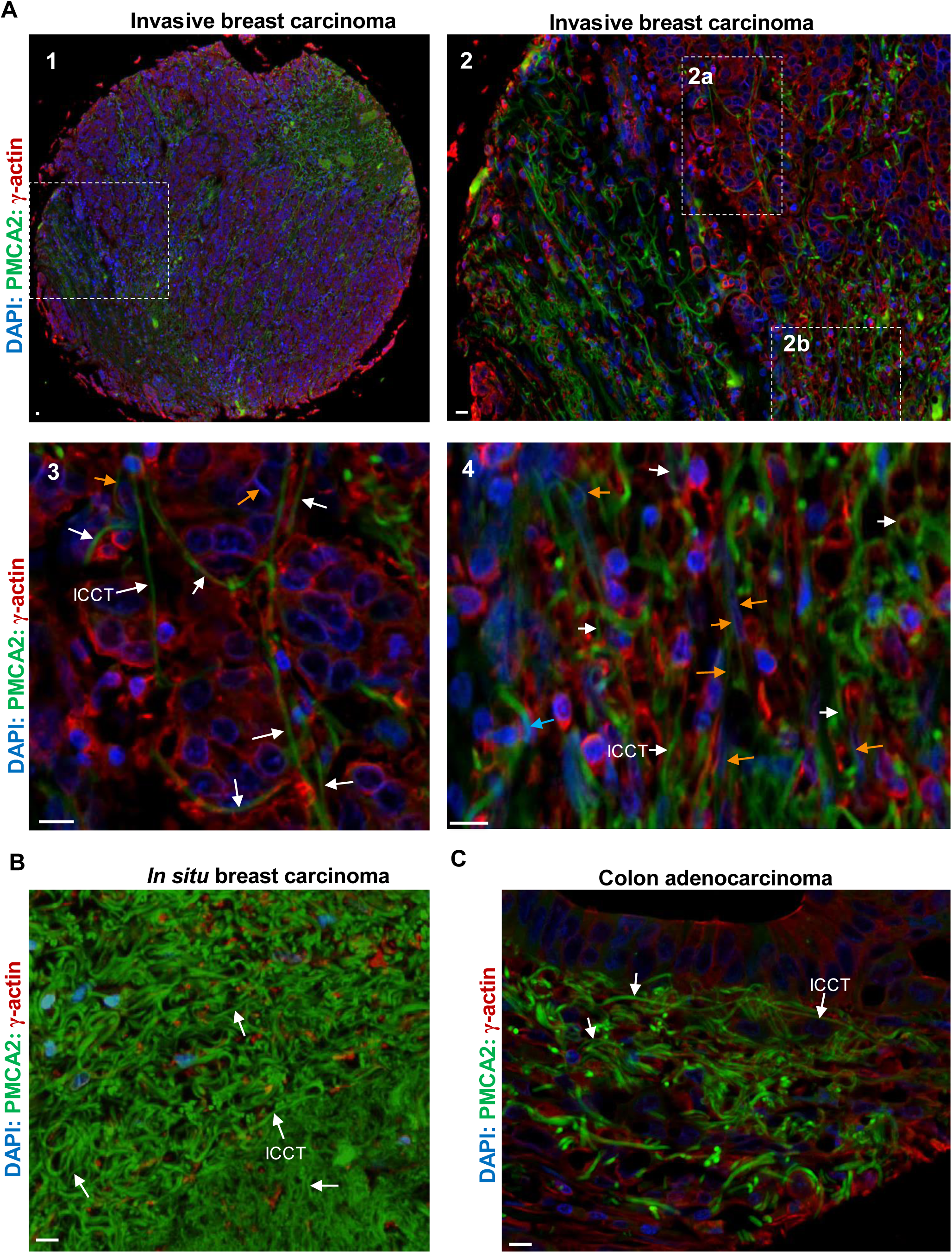
Initial CCTs present in cancer tissues *in vivo*. (**A**) Panel A1, a representative IHC fluorescence microscope image of invasive breast cancer post nuperphase tumor with many regenerated initial CCTs. The framed area is enlarged and shown in panel A2. Panel A2, there are many long or short, straight or highly-curved initial CCTs are regenerated in this area. The framed two areas of 2a and 2b are enlarged and show in the panels A3 and A4, respectively. Panel A3, there are several long, straight or curved, regenerated initial CCTs (ICCTs, white arrows), and some breast cancer cells are in migration in ICCTs (orange arrows). Panel 4, there are many regenerated initial CCTs (ICCTs, white arrows), and some breast cancer cells are in migration in ICCTs (orange arrows). There are a few established cytocapsular tubes (CCTs, cyan arrow). (**B**) In a representative image of in situ breast carcinoma, there already are large quantities of initial CCTs in high ICCT (white arrows) density. (**C**) In a representative image of clinical colon carcinoma, there are some ICCs (white arrows) in colon cancer tissues. Scale bar, 10μm.

**Extended data Fig.4.**
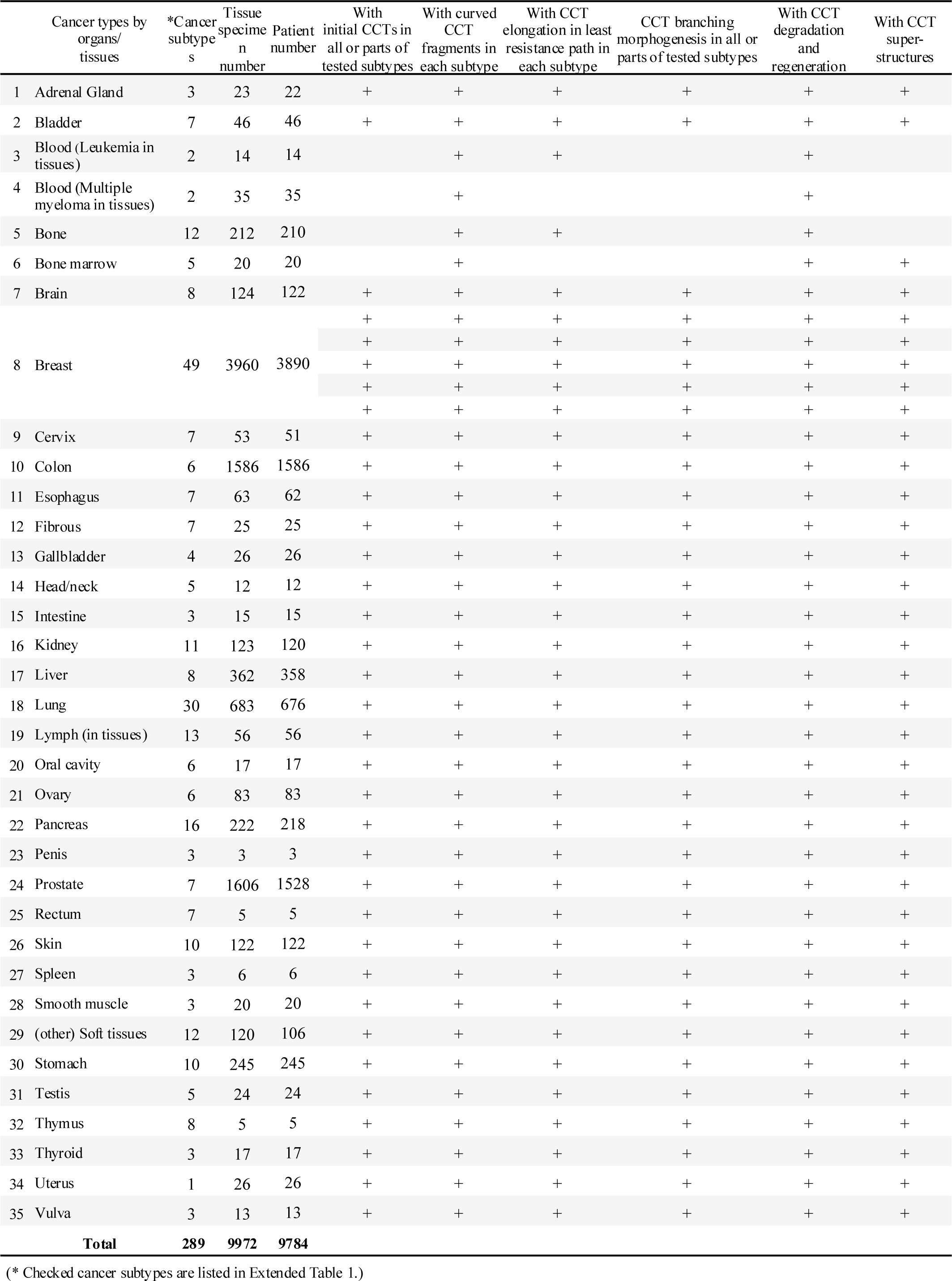
Characterization of initial cytocapsular tubes, curved cytocapsular tube fragments, CCT branching morphogenesis, CCT regeneration, and CCT superstructures in 289 types/subtypes of cancers in human organs and tissues.

**Extended data Fig.5.**
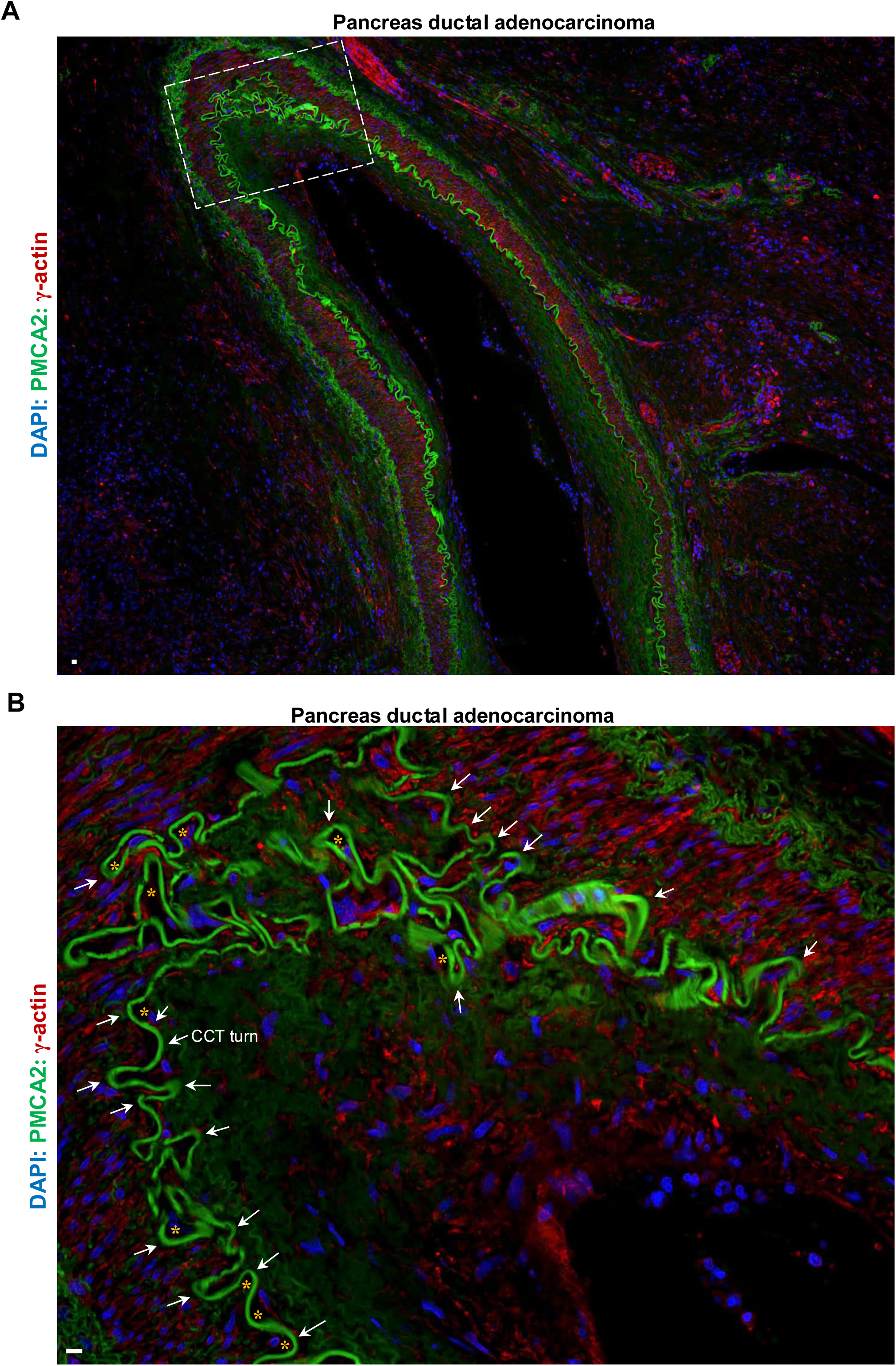
Single super-long pancreas cytocapsular tubes elongate along the path with least resistance in heterogeneous pancreas tissues and form superstructures with many curves *in vivo*. **(A)** Representative IHC fluorescence microscope image of a single pancreas cytocapsular oncocell generated, single, super-long CCT with many short/middle length curves and super-large CCT superstructure in a superlarge bent morphology in heterogeneous pancreas ductal adenocarcinoma tissue in the same plane. The macro resistances from the above and lower dense tissues force the super-long CCT to form a superlarge bent morphology. The framed area is enlarged and shown in (B). (**B**) An enlarged CCT fragment from (A). This CCT fragment has many irregular-shaped, short/middle length curves. Each pancreas CCT curve turn (white arrows) is caused by the local micro resistances from the dense tissues along the pancreas CCT elongation direction. Then, the pancreas CCT nanoprotrusion (NP) with least resistance enlarge and form enlarged NP, followed by pancreas initial CCT generation in the least resistance, and established pancreas CCT formation in the least resistance. Therefore, form a pancreas CCT turn. The orange asterisks show less or no tissue (with less or no tissue resistance) below the CCT turns. Scale bar, 10μm.

**Extended data Fig.6.**
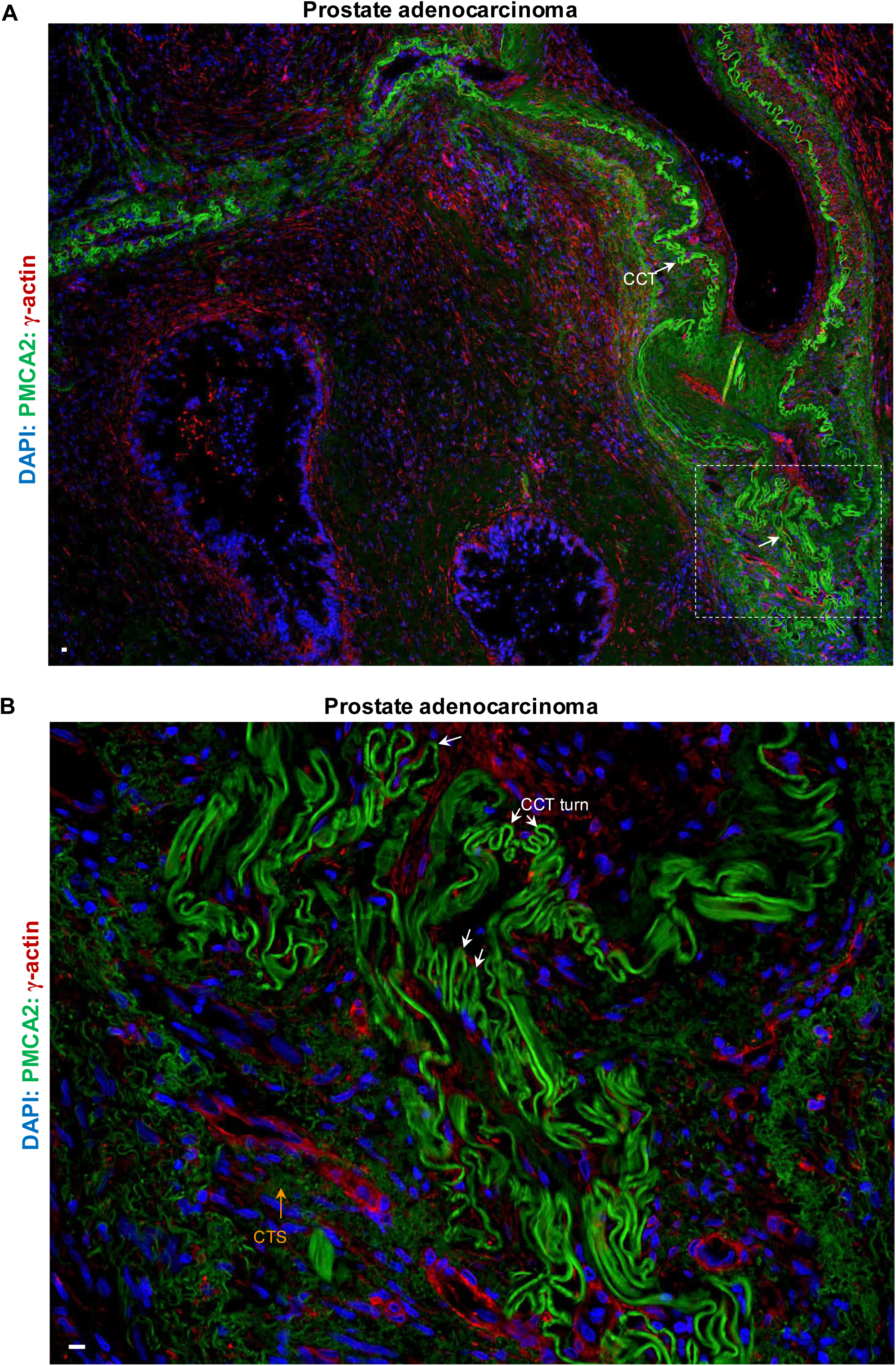
Single super-long prostate cytocapsular tubes elongate along the path with least resistance in heterogeneous prostate tissues and form superstructures with many long curves *in vivo*. **(A)** Representative IHC fluorescence microscope image of a single prostate cytocapsular oncocell generated, single, super-long CCT with many short and long curves and super-large CCT superstructure in a superlarge and irregular bent morphology in heterogeneous prostate adenocarcinoma tissue in the same plane. The macro resistances from the above and lower dense tissues force the super-long CCT to form a bent superlarge morphology. The framed area is enlarged and shown in (B). CCT (white arrows) are shown. (**B**) An enlarged CCT fragment from (A). This CCT fragment has many irregular-shaped, long curves. Each prostate CCT curve turn (white arrows) is caused by the local micro resistances from the dense tissues along the prostate CCT elongation direction. Then, the prostate CCT nanoprotrusion (NP) with least resistance enlarge and form prostate enlarged NP, followed by prostate initial CCT generation in the least resistance, and established prostate CCT formation in the least resistance. Therefore, form a prostate CCT turn. The orange asterisks show less or no tissue (with less or no tissue resistance) below the CCT turns. CCT turns (white arrows), and CCT strand (CTS, orange arrows) are shown. Scale bar, 10μm.

**Extended data Fig.7.**
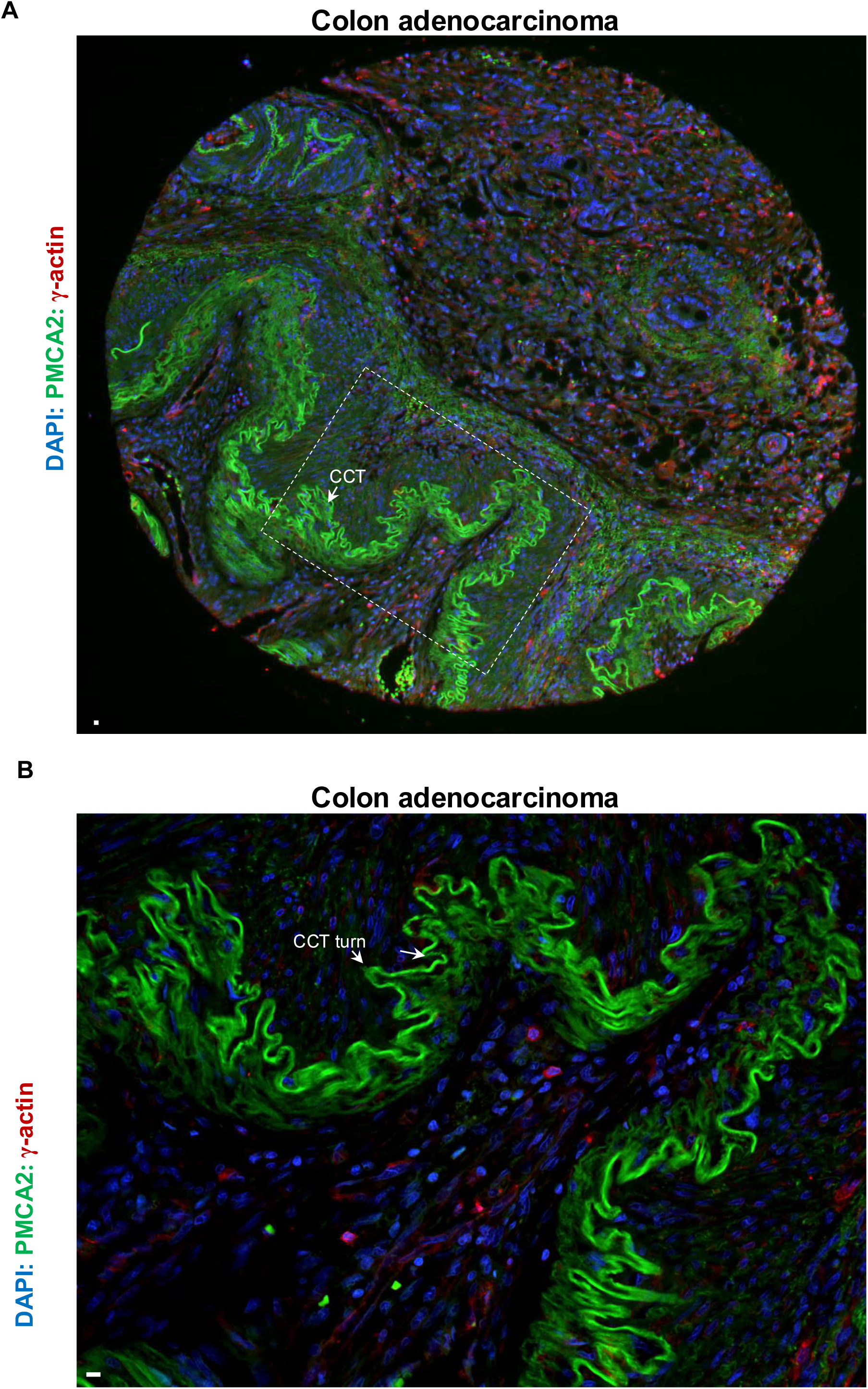
Single super-long colon cytocapsular tubes elongate along the path with least resistance in heterogeneous colon tissues and form superstructures with many curves *in vivo*. **(A)** Representative IHC fluorescence microscope image of a single colon cytocapsular oncocell generated, single, super-long colon CCT with many middle-length curves and super-large CCT superstructure in a superlarge and irregular waved morphology in heterogeneous colon adenocarcinoma tissue in the same plane. The macro resistances from the above and lower dense tissues force the super-long colon CCT to form highly-curved superlarge morphologies. The framed area is enlarged and shown in (B). CCT (white arrows) are shown. (**B**) An enlarged colon CCT fragment from (A). This colon CCT fragment has many irregular-shaped, middle-length curves. Each colon CCT curve turn (white arrows) is caused by the local micro resistances from the dense tissues along the colon CCT elongation direction. Then, the colon CCT nanoprotrusion (NP) with least resistance enlarge and form colon enlarged NP, followed by colon initial CCT generation in the least resistance, and established colon CCT formation in the least resistance. Therefore, form a colon CCT turn. CCT turns (white arrows) are shown. Scale bar, 10μm.

**Extended data Fig.8.**
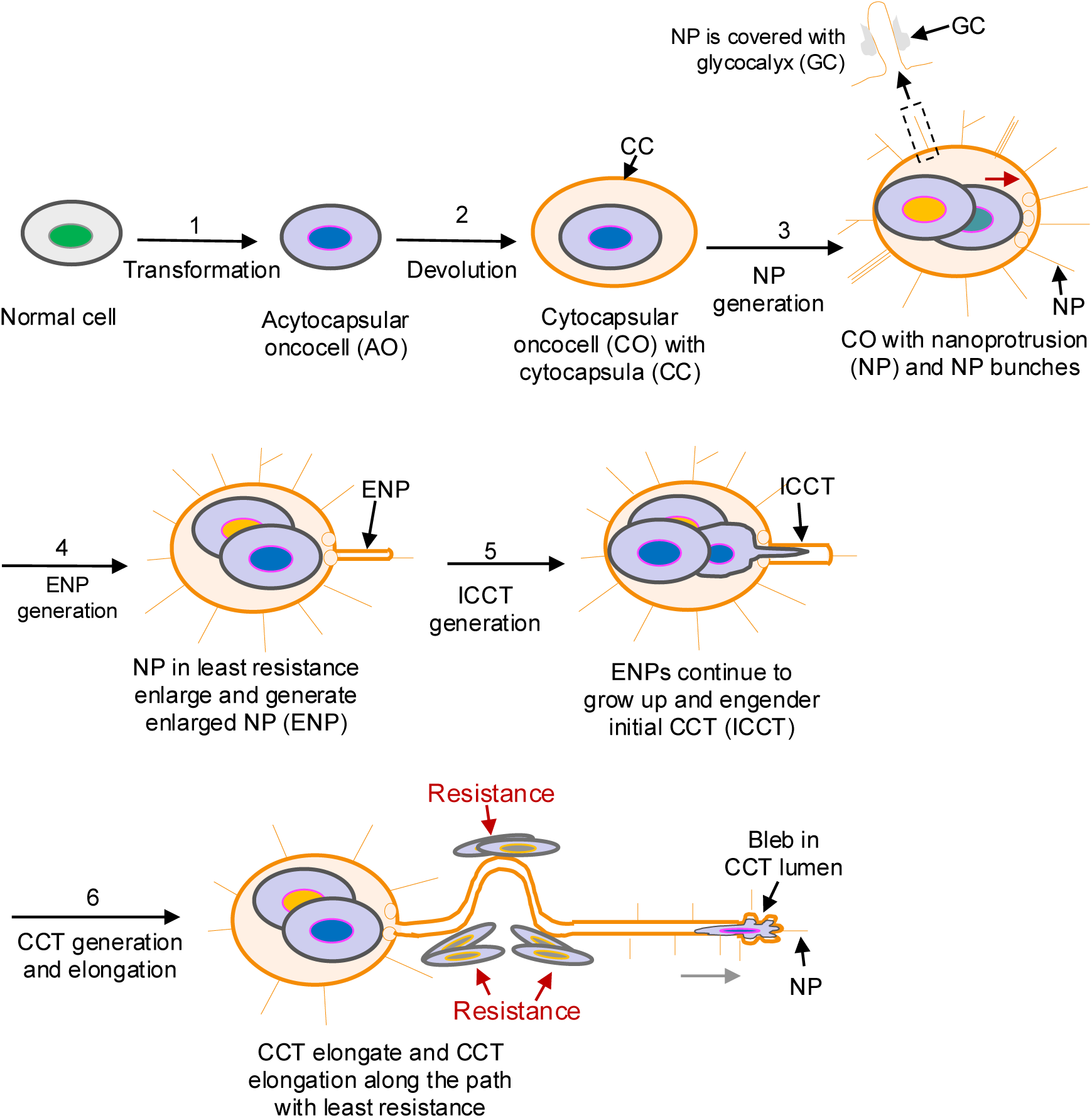
Schematic diagram of nanoprotrusion enlargement conducted cytocapsular tube elongate along the path with least resistance. There are 6 successive major steps in cytocapsular tube elongation in heterogeneous tissues *in vivo*: (**1**) Transformation: normal cell transformation generates acytocapsular oncocell (CO); (**2**) Devolution: acytocapsular oncocell devolution engenders cytocapsular oncocell with extracellular cytocapsula (CC) wrapping AO inside. The CC isolates AO from external stressful or toxic microenvironments and help AO for survival; (**3**) Nanoprotrusion (NP) generation: cytocapsular oncocells (COs) generate many nanoprotrusions (NPs, 21nm ∼ 82nm in diameter/width, 300-2700nm in length), NP bunches (40nm ∼ 152nm in width), which interconnect and form NP networks surrounding the CO. Intracytocapsular oncocells proliferate in CCs and generate cytocapsular tumors (CTs). CTs generate NP networks surrounding CTs. NPs are coated with glycocalyx in variable thickness; (**4**) Enlarged nanoprotrusion (ENP) generation: the NPs in least resistance enlarge and generate enlarged NPs (ENPs, 82nm ∼ 0.1μm); (**5**) Initial CCT (ICCT) generation: the ENPs continue to growth in size and length and generate initial CCTs (ICCTs, 0.1μm ∼ 0.5μm in diameter/width). (**6**) CCT generation and elongation: Single intracytocapsular oncocell squeezes into ICCTs, and promote ICCTs to continue to grow up in sizes and length and engender established CCTs (>0.5μm in diameter/width), therefore, CCT is elongated. CCT elongation is conducted by CCT nanoprotrusion enlargement and along the path with least resistance.

**Extended data Fig.9.**
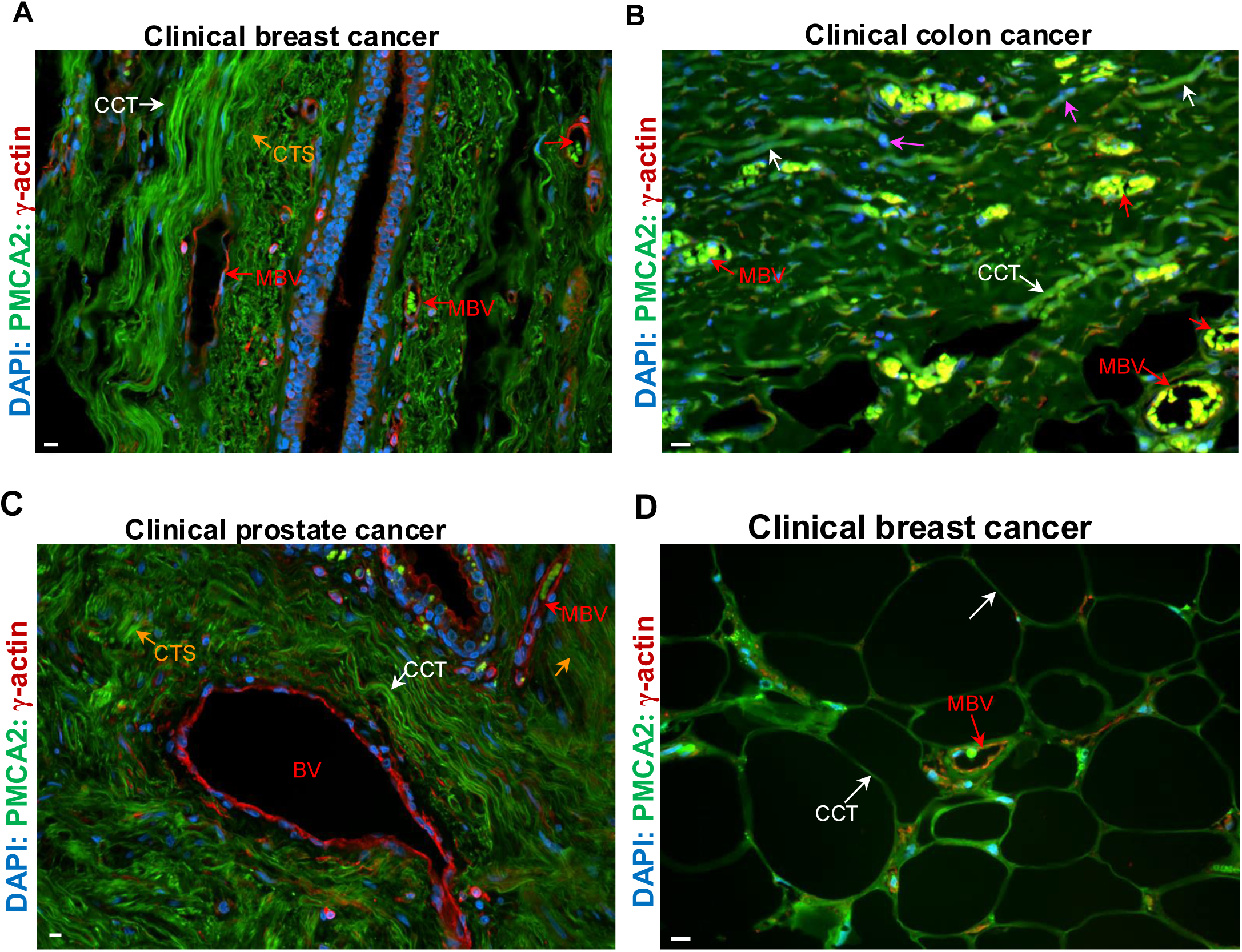
Cytocapsular tube elongate in the least resistance path lead to CCT conducted cancer metastasis occurrence much prior to CCT invasion into blood vessel and oncocell release into blood. **(A)** Representative image of clinical breast cancer tissues with many CCTs (white arrows) in the late CCT lifecycle stage of CCT degradation into CCT strands (CTSs, orange arrows). Most of breast acytocapsular oncocells (AOs) invade into breast CCTs and leave away, and there are only a few AOs left in this area. There are multiple micro blood vessels (MBVs, red arrows) in intact status without CCT invasion caused oncocell release into the blood in this area. (**B**) Representative image of clinical colon cancer tissues with many colon CCTs (white arrows) with many colon oncocells in migration in CCTs (purple arrows). Most of colon acytocapsular oncocells (AOs) invade into colon CCTs and leave away, and there are almost no colon AOs left in this area. There are multiple micro blood vessels (MBVs, red arrows) in intact status without CCT invasion caused oncocell release into the blood in this area. (**C**) Representative image of clinical prostate cancer tissues with many prostate CCTs (white arrows) in the late CCT lifecycle stage of CCT degradation into CCT strands (CTSs, orange arrows). Most of prostate acytocapsular oncocells (AOs) invade into prostate CCTs and leave away, and there are almost no prostate AOs left in this area. There are blood vessels (BV) and micro blood vessel (MBV, red arrow) in intact status without CCT invasion caused oncocell release into the blood in this area. **(D)** Representative image of clinical breast cancer tissues with large CCTs (white arrows) networks in the late stage. Most of breast acytocapsular oncocells (AOs) invade into breast CCT networks and leave away, and there is no breast AOs left in this area. There are a micro blood vessel (MBV, red arrows) in intact status without CCT invasion caused oncocell release into the blood in this area. Scale bar, 10μm.

**Extended data Fig.10.**
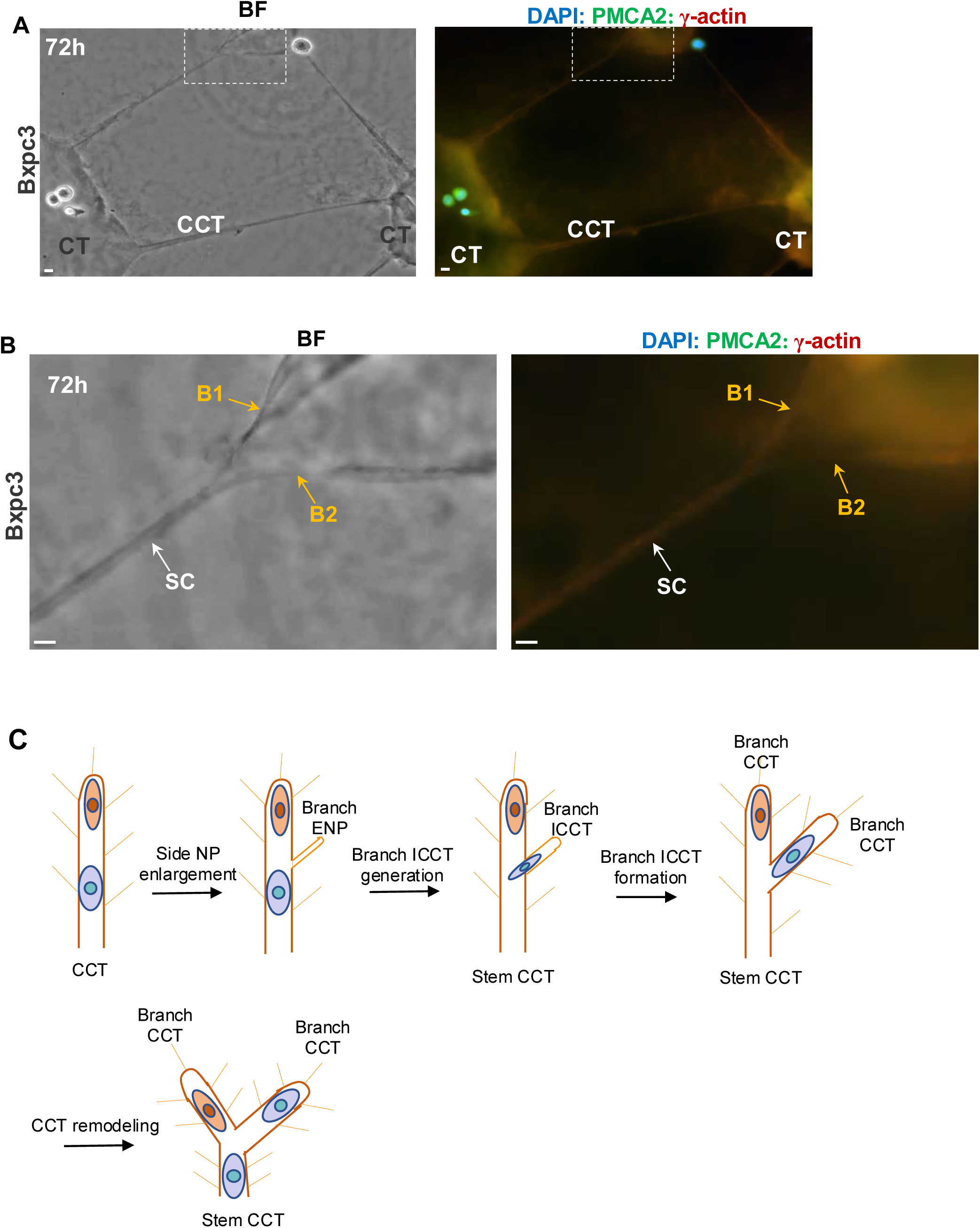
CCT branching morphogenesis in side NP enlargement conducted bifid branching morphogenesis format in vitro. (**A**) Representative bright field (BF) and fluorescence microscope images of CCT branching morphogenesis CC/CCT culture kits *in vitro*. At 72h after Bxpc3 pancreas cancer cell implantation in CC/CCT culture kits, pancreas cytocapsular tumorspheres (CTs), multiple CCTs interconnect CTs, and CCT branches are generated. The framed area in BF and fluorescence microscope images are enlarged and shown in (B). Scale bar, 10μm. (**B**) Enlarged BF and fluorescence microscope images of framed areas in (A). The stem CCT (SC) side nanoprotrusions in least resistance generate enlarged NPs, followed by initial CCT (ICCT) and established CCT generation, and engender a branch CCT. After CCT membrane remodeling, the branch CCT and stem CCT are remodeled into two near similar CCT branches (branch 1 (B1) and branch 2 (B2), orange arrows). This kind of CCT branching morphogenesis format is named as bifid CCT branching morphogenesis format. Scale bar, 10μm. (**C**) Schematic diagram of CCT bifid branching morphogenesis format, which contains 4 successive major steps: (1) Side NP enlargement: CCT side NPs in least resistance enlarge and generate branch enlarged NPs (branch ENPs); (2) Branch initial CCT (ICCT) generation: Branch ENPs continue to grow up into branch ICCT; (3) Branch CCT formation: With intracytocapsular oncocell riven ICCT growth in size and length, branch ICCTs continue to grow up into branch CCTs. (4) CCT remodeling: the connected cytocapsular membranes of stem CCT and branch CCT are remodeled into two similar or near similar CCTs (in diameter/width) and interconnecting with the stem CCT, and form two branch CCTs.

**Extended data Fig.11.**
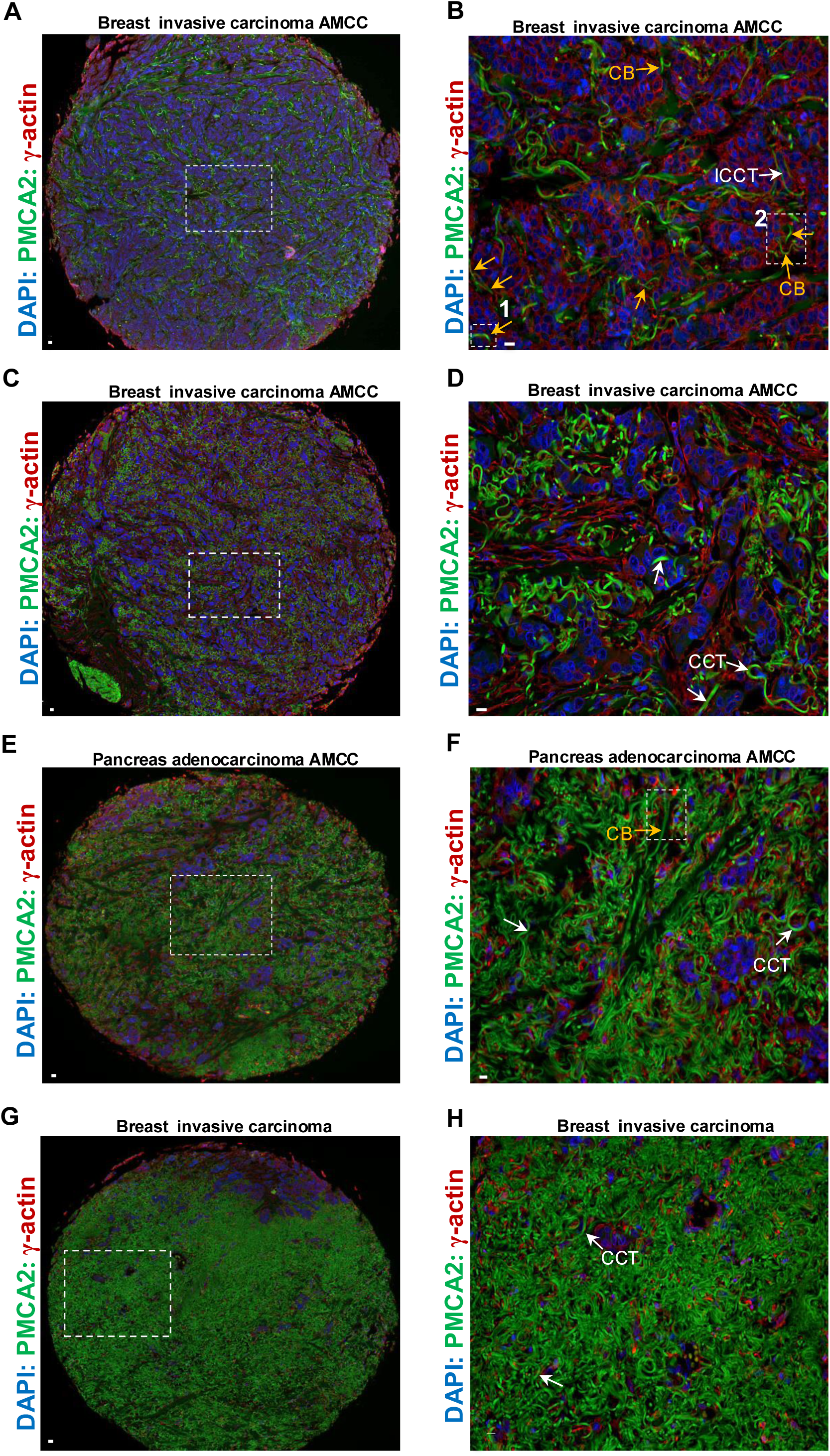
CCT bifid branching morphogenesis significantly increase CCT numbers and promote cancer metastasis in tissues *in vivo*. (**A**) Representative IHC fluorescence image of breast invasive carcinoma in the early CCT regeneration stage I in acytocapsular oncocell mass-CC/CCT complex (AMCC stage I). The framed area is enlarged and shown in (B). (**B**) Enlarged framed area in (A). In the dense acytocapsular oncocell mass, regenerated initial CCTs (ICCTs) and CCTs perform bifid branching morphogenesis and increase CCT numbers. In AMCC stage I, ICCTs/CCTs occupied areas are 0-10% in areas in the AMCC. Multiple CCT branches (CBs, orange arrows) and ICCTs (white arrows) are shown. The two framed areas (area 1 and area 2) with CCT branches are enlarged and shown in Figure 2A1 and 2A2 panels, respectively. (**C**) Representative IHC fluorescence image of breast invasive carcinoma in AMCC stage II. The framed area is enlarged and shown in (D). (**D**) Enlarged framed area in (C). In AMCC stage II, ICCTs/CCTs/CCT strands occupied areas are 10%-40% in areas in the AMCC. (**E**) Representative IHC fluorescence image pancreas adenocarcinoma in the AMCC stage III. The framed area is enlarged and shown in (F). (**F**) Enlarged framed area in (E). In AMCC stage III, ICCTs/CCTs/CCT strands occupied areas are 40%-70% in areas in the AMCC III. The framed area with CCT branch is enlarged and shown in Fig2A3 panel. (**G**) Representative IHC fluorescence image of breast invasive carcinoma in the AMCC stage IV. The framed area is enlarged and shown in (H). (**H**) Enlarged framed area in (G). In AMCC stage IV, ICCTs/CCTs/CCT strands occupied areas are 70%-100% in areas in the AMCC III. CCTs (white arrows) are shown. With CCT number significantly increase (from A to C, to E, to G, from AMCC stage I to stage II, stage III, and stage IV), more and more acytocapsular oncocells (AOs) invade into CCTs via alloentry, and leave away, and less and less AOs left. Scale bar, 10μm.

**Extended data Fig.12.**
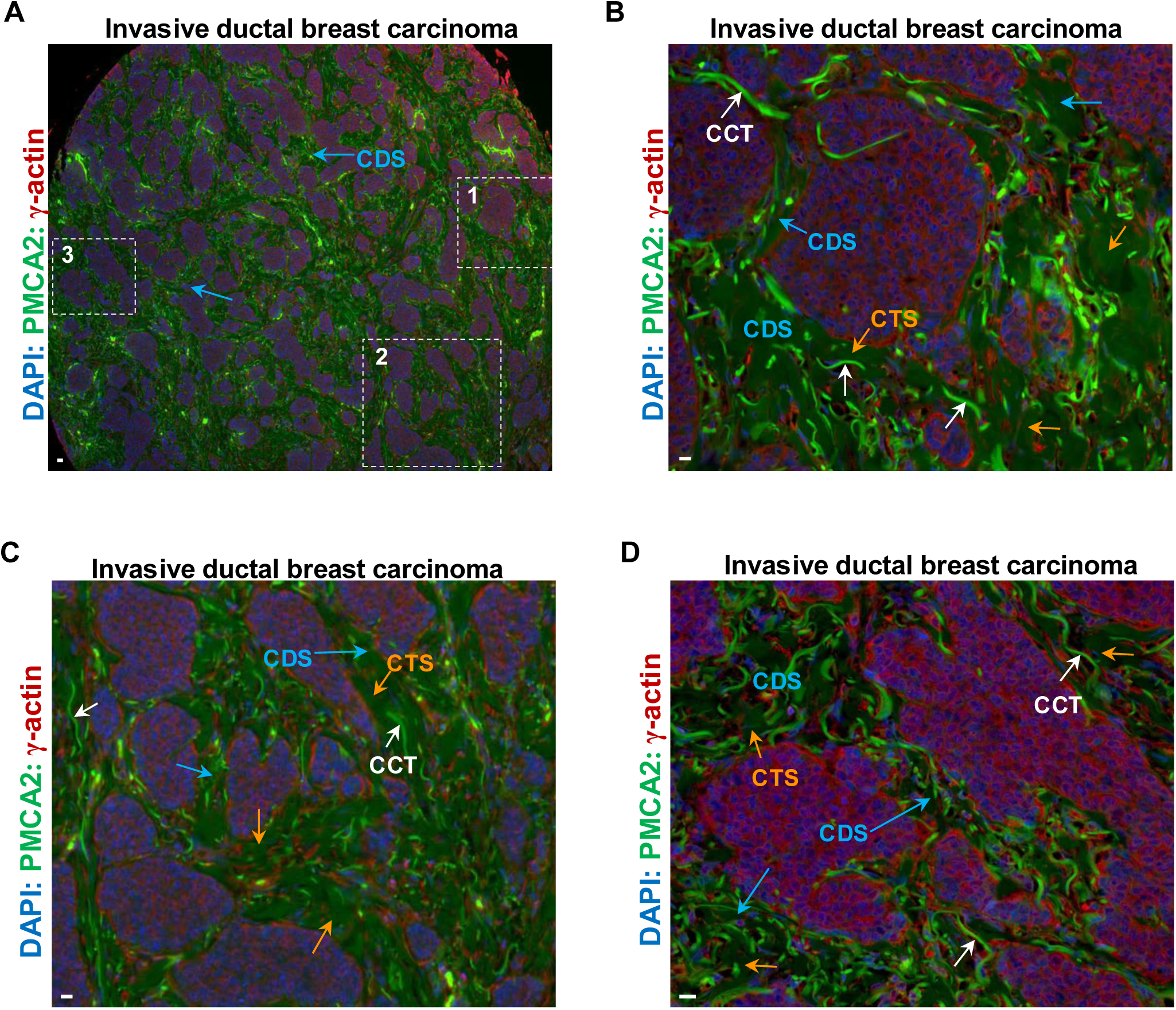
CCT regeneration associates with new cycle of cancer metastasis in AMCC. (**A**) Representative invasive ductal breast carcinoma in AMCC with many CCT degradation left cavity strips (CDSs) interconnected CDS networks. There are many CCT degradation left cavity strips (CDSs) in variable widths crossing in the dense acytocapsular oncocell masses, and separate dense acytocapsular oncocell (AO) mass into many dense AO mass islands. The CCTs degrade into cloud-like CCT strands (smear cloud in green color in the image) followed by disappearance. The framed areas (1, 2, and 3) are enlarged and shown in the panel (B, C, D), respectively. (**B**) Enlarged area 1 in (A). Many new CCTs are regenerated and elongate in the previous CCT degradation left cavity strips (CDSs). Most regenerated CCTs elongate in CDSs with less resistances, a few CCTs invade into dense AO mass islands. (**C**) Enlarged area 2 in (A). There are multiple CDSs interconnect and form CDS network for regenerated new CCTs to elongate. (**D**) Enlarged area 2 in (A). In the CDSs, there are many regenerated new CCTs, which tightly contact with dense AO islands, and allow AOs invade into CCTs via alloentry, followed by new cycles of cancer metastasis in AMCC. Scale bar, 10μm.

**Extended data Fig.13.**
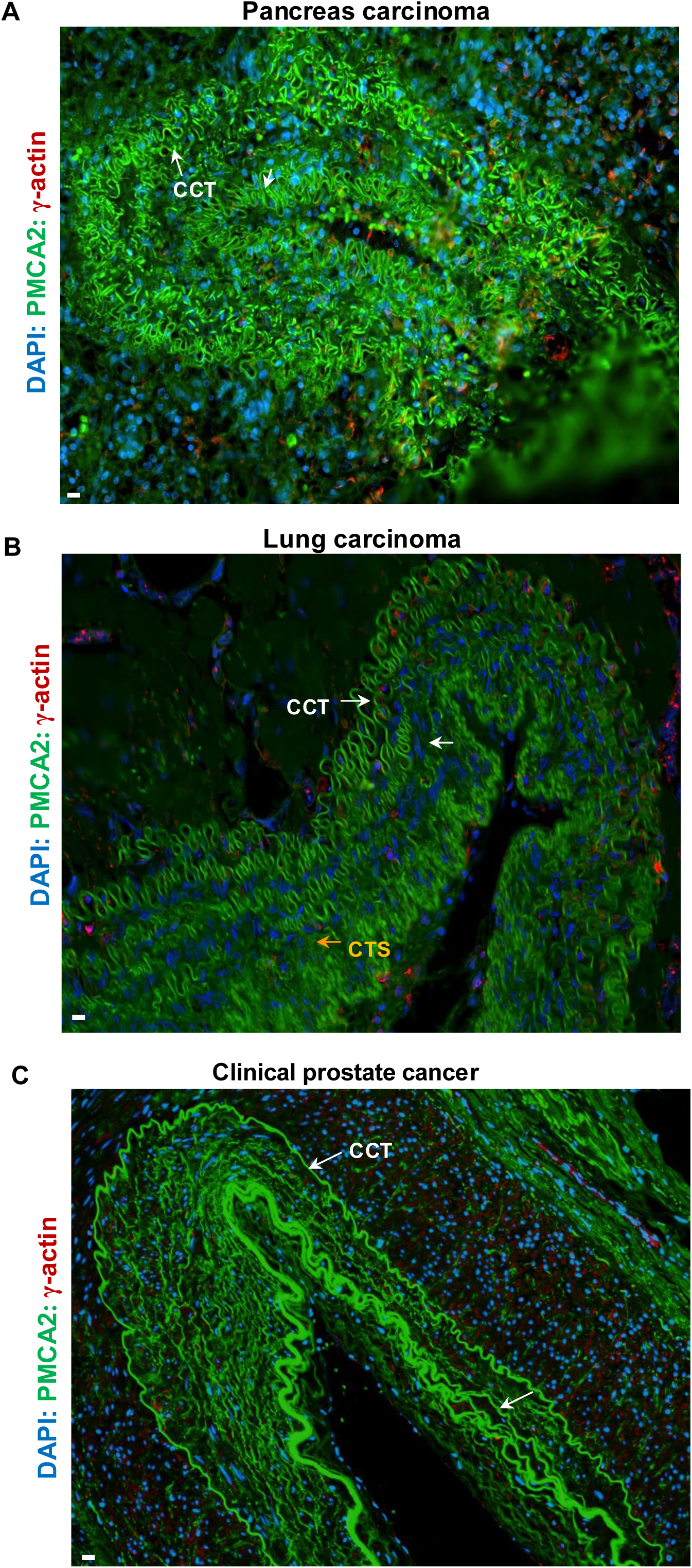
CCT mega superstructures in pancreas, lung and prostate cancers in vivo. (**A**) Representative IHC fluorescence image of a part of a super-large pancreas CCT mega superstructures with three-layered, irregular triangle-shaped, and highly curved CCT superstructures with many CCT curves/coils of each CCT superstructure. (**B**) Representative image of a super-large lung CCT mega superstructures with five-layered, irregular-shaped CCT superstructures with many CCT curves/coils of each CCT superstructure. Some CCTs are in degradation into CCT strands (CTSs, orange arrows). (**C**) Representative image of a super-large prostate CCT mega superstructures with more than seven-layered, irregular-shaped CCT superstructures with many CCT curves of each CCT superstructure.

## Notes

### Competing Interest Statement

The authors have declared no competing interest.

### Summary of Updates

"for metastasis initiation" is added in the title; some verbal modifications are added in the figure legends; labels are added in Figs. 1, 2, 5, Extended Data figures 1, 2, 9; a new Fig.5 is added in the main figures.

